# Competition-based screening secures the evolutionary stability of a defensive microbiome

**DOI:** 10.1101/2020.11.24.395384

**Authors:** Sarah F. Worsley, Tabitha M. Innocent, Neil A. Holmes, Mahmoud M. Al-Bassam, Barrie Wilkinson, J. Colin Murrell, Jacobus J. Boomsma, Douglas W. Yu, Matthew I. Hutchings

**Author notes:** These authors contributed equally to the manuscript (Tabitha M. Innocent, Neil A. Holmes).

## Abstract

Cuticular microbiomes of *Acromyrmex* leaf-cutting ants are exceptional because they are freely colonizable, and yet the prevalence of *Pseudonocardia*, a native vertically transmitted symbiont that controls *Escovopsis* fungus-garden disease, is never compromised. Game theory suggests that *competition-based screening* can allow the selective recruitment of antibiotic-producing bacteria from the environment, by fomenting and biasing competition for abundant host resources. Mutual symbiont aggression benefits the host and also maintains native symbiont viability. Here we use RNA-stable isotope probing (RNA-SIP) to confirm predictions that *Acromyrmex* cuticles can maintain a range of microbial symbionts. We then used dual-RNA-sequencing and bioassays to show that vertically transmitted *Pseudonocardia* strains produce antibacterials that differentially reduce the growth rates of other microbes, ultimately eliminating non-antibiotic-producing strains that might parasitize the symbiosis while still allowing antibiotic-producing *Streptomyces* strains to survive. Open cuticular microbiomes can thus maintain a specific co-evolved mutualism by restricting access for other bacterial strains.

## Introduction

The diversity of insect associated microbial communities is staggering. They may consist of single intracellular symbionts with reduced genomes owing to coadaptation at one extreme^1^, to dynamic microbiomes in open host compartments such as guts at the other end of the scale^2^. Gut microbiomes have been most intensively studied in humans and other vertebrates because there is increasing consensus of their vital interest for host fitness throughout ontogenetic development^3, 4, 5^. The stability and cooperative characteristics of complex microbiomes is a paradox. While relentless competition is the default setting of the microbial world^6^, hosts appear to evolve control by holding their microbiome ecosystems on a leash^7^, but how dynamic stability under continuing turnover is achieved remains unclear. Despite an abundance of microbiome research, recent reviews have concluded that “integration between theory and experiments is a crucial ‘missing link’ in current microbial ecology”^8^ and that “our ability to make predictions about these dynamic, highly complex communities is limited” ^9^.

Game theory suggests a compelling solution to the unity-in-diversity paradox by showing that *competition-based screening* can be a powerful mechanism to maintain cooperative stability. Screening is likely to work when hosts evolve to: (1) provide nutrients and/or space to foment competition amongst symbionts, thus creating an attractive but demanding environment, and (2) skew the competition so that the mutualistic symbionts enjoy a comparative advantage. Competitive exclusion then ‘screens in’ mutualists and ‘screens out’ parasitic or free-rider symbionts^10, 11, 12^. Screening is conceptually clearest when the symbiont trait that confers competitive superiority is the same as (or strongly correlated with) the trait that benefits the host. An illustrative example of such correlated functionality was provided by Heil^13^ who showed that ant-hosting acacia plants provide copious food bodies, which fuels the production of numerous externally patrolling ant workers. The ant species whose colonies invest in a greater number of, and more active, workers typically wins the plant, and the same investment in active workers is likely to better protect the host plants against herbivores^13^.

Screening has also been suggested to act in animal-microbe symbioses. For instance, Tragust et al.^14^ showed that carpenter ants acidify their own stomachs by swallowing acidopore secretions. Entomopathogenic bacteria are rapidly killed off, whereas the gut bacterial symbiont *Asiaia* sp. (Acetobacteraceae) exhibits a lower mortality rate and maintains itself in the midgut. Addressing a similar question, Itoh *et al*.^15^ used co-inoculation experiments to show that environmentally recruited but co-adapted ‘native’ *Burkholderia* symbionts outcompete non-native bacteria in the gut of their bean bug host, even though they are able to establish in the absence of the ‘native’ symbiont. Finally, Ranger *et al*.^16^ showed that ambrosia beetles selectively colonize physiologically stressed trees, which have a high ethanol titre due to anaerobic respiration. The vertically transmitted fungal symbionts of these beetles have evolved to detoxify the ethanol whereas competing weedy fungi remain inhibited.

Competition-based screening seems particularly apt for the establishment of protective microbiomes when competition would screen in microbes that are similarly able to produce compounds that kill competitors and thus contribute to defensive functions from the host’s perspective^7, 10, 11, 17^. Moreover, natural selection is expected to reinforce the correlation between antibiotic production and antibiotic resistance, since production without resistance would be suicidal. Antibiotic-resistant strains should thus be superior competitors in antibiotic-filled environments, reinforcing the abundance of antibiotic producers as well^11^. Previous research has shown that the protective, cuticular microbiome of *Acromyrmex echinatior* leafcutter ants (Formicidae, Attini) is an ideal model system to test the dynamic predictions of screening theory in a context that also acknowledges the long-term ecosystem-on-a-leash perspective^7, 18^. These ants forage for fresh leaf fragments to provision their co-evolved fungus-garden mutualist *Leucoagaricus gongylophorus*^19, 20^. The fungal cultivar produces gongylidia, nutrient-rich swellings that are the sole food source for the queen and larvae^21, 22^ and the predominant food source for the workers who only ingest plant-sap in addition to fungal food^23^. However, *Leucoagaricus* is at risk of being parasitized by the specialized, coevolved mould *Escovopsis weberi,* which can degrade the fungal cultivar and also cause severe ant paralysis and mortality^24, 25, 26, 27^. To prevent infections, leafcutter ants have evolved a range of weeding and grooming behaviours^28, 29, 30^, and *A. echinatior* and other *Acromyrmex* species also maintain filamentous actinomycete bacteria that grow as a white bloom on the cuticles of large workers and produce antimicrobials that inhibit the growth of *E. weberi* ^26^. In Panama, where almost all fieldwork on this multipartite symbiosis has been carried out, the cuticle of *Acromyrmex* workers is dominated by one of two vertically transmitted strains of *Pseudonocardia*, named *P. octospinosus* (Ps1) and *P. echinatior* (Ps2)^18, 31, 32, 33^.

Newly eclosed large workers are inoculated with the vertically transmitted *Pseudonocardia* strain by their nestmates upon emergence. This blooms over the cuticle, reaching maximum coverage after ca. six weeks, before shrinking back to the propleural plates as the ants mature to assume foraging tasks^34, 35^. Several studies have also identified other actinomycete strains on the propleural plates of *A. echinatior*, which are presumed to be acquired from the environment^17, 18, 36, 37, 38, 39^. This includes species of the bacterial genus *Streptomyces*, which have been identified on ants maintained in laboratory-based colonies ^36, 37, 38^, as well as in 16S rRNA gene amplicon sequencing studies of ants sampled from their native environment, although many of these could also have been close relatives of *Streptomyces*^18^. Species of this actinomycete genus produce a variety of antimicrobials so their additional presence may suggest a form of multi-drug therapy against *Escovopsis*^37, 38, 39^. However, these putative functions remain enigmatic because *Streptomyces* symbionts were never found on the callow workers^18^ that execute hygienic and defensive fungus-garden tasks^35^. In terms of resources, the propleural plates have a high concentration of tiny subcuticular glands, which are presumed to supply the cuticular microbiome with resources^40^. These plates can thus be conjectured to create the food-rich but antibiotic-laden demanding environment that competition-based screening assumes, because the vertically transmitted native *Pseudonocardia* symbiont always colonizes the propleural plates first^11^. We have previously shown that both *P. octospinosus* and *P. echinatior* encode and make antibacterial compounds that inhibit multiple unicellular bacteria, but not *Streptomyces* species^31^. However, the other key elements of the screening hypothesis have remained untested.

The present study provides a number of tests of competition-based screening, using the cuticular microbiome of *A. echinatior* as an open symbiotic ecosystem model where the host holds the resource leash. Within these constraints, the single co-adapted and dominant defensive *Pseudonocardia* symbiont competes with an unspecified number of additional symbionts, which have unknown functions but are predicted by screening theory to keep the microbiome cooperative by excluding parasitic or free-riding microbes. We used RNA-SIP (stable isotope probing) to show that the ants provide a public food resource on their cuticles that is indeed consumed by multiple bacterial species. We then use dual RNA sequencing to show, *in vivo,* that mutualistic *P. octospinosus* and *P. echinatior* strains express antibacterial biosynthetic gene clusters (BGCs) on the ant cuticle. Next, we show that diffusible metabolites of these *Pseudonocardia* species exhibit broad-spectrum antibacterial activity *in vitro*, but only weakly inhibit *Streptomyces* species, which we separately and directly show are resistant to a range of antibiotics. Finally, we use *in-vitro* competition experiments to show that slower-growing *Streptomyces* species can competitively exclude faster-growing non-antibacterial-producing species, but only when grown on media infused with *Pseudonocardia* metabolites. Together, these results let us conclude that competition-based screening is a plausible mechanism for the environmental acquisition of additional strains with similar mutualistic inclinations as the native symbiont, and that screening thus maintains and dynamically stabilizes the defensive microbiome initiated by the single native *Pseudonocardia* symbiont. Our results underline that a host leash of a symbiotic microbiome can be restricted to a single co-adapted symbiont that further shapes the microbiome to its own and the host’s interests, provided its metabolites are hostile enough to avoid exploitation by non-beneficial invaders.

## Results

### The host provides public resources to its cuticular microbiome

RNA-SIP tracks the flow of heavy isotopes from the host to the RNA of microbial partners that metabolize host-derived resources^41, 42, 43^. Labelled and unlabelled RNA within a sample are separated and used as templates for 16S rRNA amplicon sequencing, so that the bacterial taxa that do (and do not) use host-supplied resources can be identified^43^. Three replicate groups of 22 mature worker ants were fed a 20% (w/v) solution of either ^12^C or ^13^C glucose for 10 days and then propleural plates were dissected out for total RNA extraction (Supplementary Figure 1). Control feeding experiments demonstrated that a fluorescently labelled glucose-water diet was not transferred to the surface of the propleural plate region of the ants during feeding, and the ants were only ever observed to feed using their mandibles such that their propleural plates never came into direct contact with the liquid diet (Supplementary Figure 2).

Following cDNA synthesis, 16S rRNA gene amplicon sequencing showed that filamentous actinomycetes dominate the propleural plate samples, making up 76.6 % and 78.0 % of unfractionated cDNA samples from ^12^C and ^13^C fed ants, respectively (Figure 1). The most abundant bacterial genera were *Pseudonocardia*, 35.8% and 38.1% in ^12^C and ^13^C samples respectively, and *Streptomyces*, 19.7% and 20.5%, respectively. *Wolbachia* made up 22.8% and 19.6%, respectively. *Wolbachia* are known to be associated with the thoracic muscles of *A. echinatior* worker ants where they may have an unspecified mutualistic function^44, 45^. We therefore assume these reads came from residual ant tissue on the dissected propleural plates and do not consider them further. More than 95% of the *Pseudonocardia* 16S rRNA gene reads in unfractionated RNA samples were identical in sequence to a single *P. octospinosus* mutualist strain^18, 31^. The presence of *Streptomyces* bacteria is consistent with previous studies and supports the hypothesis that additional antibiotic-producing actinomycetes can be horizontally acquired by these large workers ^17, 36, 37, 38, 39, 46^ after always first being inoculated with the native *Pseudonocardia* symbiont^18^.

**Figure 1.**
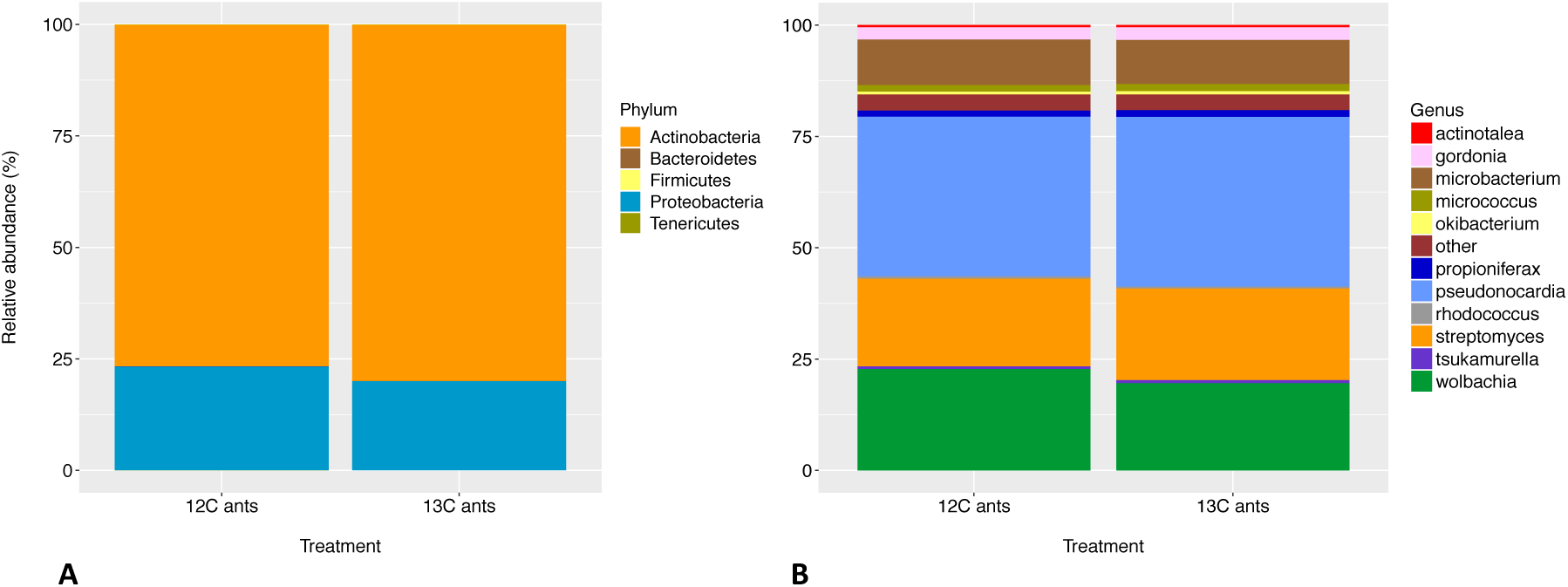
Frequencies of bacteria at different taxonomic levels in unfractionated RNA samples from the propleural plates of ants provided with a 20% (w/v) solution of either ^12^C- or ^13^C- labelled glucose. (**A**) Phylum resolution, showing that >75% were actinobacteria. The remaining reads almost all corresponded to *Wolbachia* (Proteobacteria) so the three other phyla do not appear in the bars (total relative abundance of 0.6% and 0.4% in ^12^C and ^13^C fractions, respectively). As *Wolbachia* is not part of the cuticular microbiome (see text) these results show that the cuticular microbiome is completely dominated by actinobacteria. (**B**) Genus-level resolution showing that the cuticular microbiomes are dominated by the native *Pseudonocardia* symbiont followed by appreciable fractions of horizontally acquired *Streptomyces* species and a series of other actinobacteria at low prevalence.

Caesium trifluoroacetate density gradient ultracentrifugation was used to separate the ^13^C-labelled “heavier” RNA from the un-labelled ^12^C “lighter” RNA within the ^13^C-fed samples (Supplementary Figure 1). RNA samples from the control ^12^C dietary treatment also underwent density gradient ultracentrifugation. The resulting gradients were fractionated by buoyant density (Supplementary Figures 3 & 4), and cDNA from each fraction was used in quantitative RT-PCR reactions. This demonstrated that 16S rRNA gene copies had shifted to higher buoyant densities in samples derived from the ^13^C dietary treatment, which is consistent with the heavier ^13^C isotope being incorporated into the RNA of cuticular bacteria, due to the metabolism of labelled host resources. Fractions spanning peaks across the buoyant density gradient were selected for 16S rRNA gene amplicon sequencing (Supplementary Figure 4), which confirmed that the actinobacterial genera had shifted to higher buoyant densities under the ^13^C treatment (Figure 2A and B). *Pseudonocardia* and *Streptomyces* sequences had respective relative abundances of 36.30% ± 4.17 (SE) and 19.18 ± 0.33 in fraction 7, and 32.30% ± 4.11 and 17.64 ± 0.26 in fraction 8 of the ^13^C samples (Figure 2B and C). These fractions had an average buoyant density of 1.797 g ml^-1^ ± 0.001 SE (fraction 7) and 1.805 g ml^-1^ ± 0.001 SE (fraction 8) and coincided with the peak in 16S rRNA gene copy number identified by qPCR (Supplementary Figures 3 & 4).

**Figure 2.**
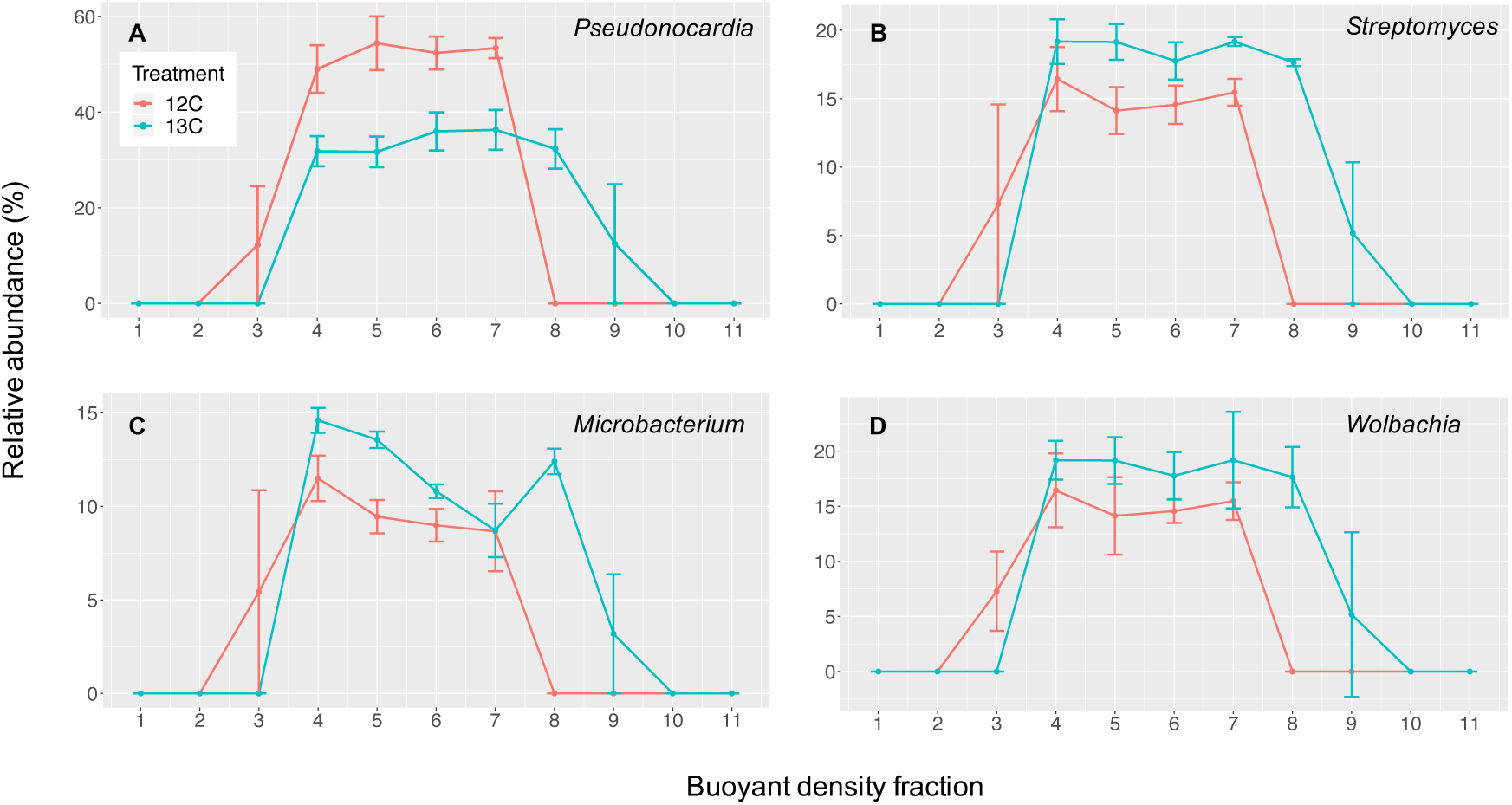
Average frequencies of the genera *Pseudonocardia* (**A**), *Streptomyces* (**B**), *Microbacterium* (**C**), and *Wolbachia* (**D**) across fractions of the buoyant density gradient of samples taken from ants fed on a ^13^C (blue) or ^12^C (red) glucose diet. Higher fraction numbers represent greater buoyant densities (g ml^-1^), and only bacteria from the ^13^C-treated ants were found in the heavier fractions on the right, indicating uptake of heavier ^13^C from the ant host. Error bars indicate standard errors with n=3 replicate samples per treatment.

In contrast, bacterial sequences from the ^12^C samples were barely detectable by qPCR in fraction 8 (buoyant density of 1.801 g ml-1 ± 0.003) (Supplementary Figure 4). *Pseudonocardia* and *Streptomyces* sequences were observed in fraction 7 of ^12^C samples (Figure 2A and B), but only 1.59% ± 1.20 of the bacterial 16S rRNA sequences could be amplified in qPCR experiments (Supplementary Figure 4). Peaks in 16S rRNA gene copy number instead occurred in fractions 5-6 of ^12^C samples, which had average buoyant densities of 1.780-1.788 g ml^-1^ (Supplementary Figures 3 & 4). The other horizontally acquired taxa showed similar shifts to higher buoyant densities between the ^12^C and ^13^C treatment groups; for example, the genus *Microbacterium* made up 8.71% ± 1.43 of fraction 7, and 12.40% ± 0.68% of fraction 8, respectively (Figure 2C). Note that all these percentages become ca. 20% higher after excluding *Wolbachia*, i.e. when considering only the cuticular microbiome.

In addition to the shifts in buoyant density, indicating uptake of ^13^C by multiple bacterial genera from the ant, several of the genera were also more abundant in the heavy fractions of the ^13^C samples (which correspond with the peak in 16S rRNA gene copy number identified by qPCR, Supplementary Figure 4) than in the heaviest fractions of the ^12^C samples (Supplementary Figure 5), implying that they were particularly effective in their use of carbon-based resources from their host ants. For example, *Streptomyces* species increased from an average frequency of 14.8% ± 0.33% (SE) in ^12^C heavy fractions to 18.5% ± 0.29% in ^13^C heavy fractions (Supplementary Figure 5). Interestingly, although *Pseudonocardia* was abundant in the heaviest fractions of the ^13^C samples, its frequency decreased from an average of 53.65% ±1.95% to 34.30 ±3.78% between the heaviest fractions of the ^12^C treatment and ^13^C treatment (Supplementary Figure 5). This corresponded with a concurrent increase in the abundance of other genera and a more even representation of genera in the heavier ^13^C fractions. These increases in the other genera (all actinobacteria) are further evidence that ant-derived resources were not solely being made available to the *Pseudonocardia* (which otherwise would have dominated the ^13^C heavy fractions) but are publicly available to all bacteria on the cuticle.

The frequency of *Wolbachia* also increased from 9.86% ± 1.60% in the heavy ^12^C fractions, to 19.49% ±3.53% in the ^13^C heavy fractions (Figure 2D; Supplementary Figure 5). Since *Wolbachia* are extracellular endosymbionts, this finding supports the interpretation that resources were supplied to cuticular bacteria by the ant hosts and not taken directly from the glucose water. This interpretation is also supported by isotope ratio mass spectrometry (IRMS), which showed that surface-washed ants incorporate a significant amount of the ^13^C from their glucose diet into their bodies (Supplementary Figure 6), and by direct fluorescent microscopy demonstrating that the glucose water was not transferred to the propleural plate (Supplementary Figure 2).

### Antibacterial BGCs are expressed by *Pseudonocardia* on the ant cuticle

We previously generated high-quality genome sequences for five *P. octospinosus* and five *P. echinatior* strains isolated from *A. echinatior* ant colonies and identified several BGCs (biosynthetic gene clusters) in each of their genomes that are associated with antimicrobial activity ^31^. To establish if these BGCs are expressed *in vivo* on the ant cuticle, total RNA was extracted and sequenced from the propleural plates of ants in the captive colonies Ae088 (which hosts a vertically transmitted *P. echinatior* strain) and Ae1083 (which hosts a *P. octospinosus* strain) (Supplementary Table 1). A single RNA extraction was carried out for each colony, with each sample consisting of the pooled propleural plates of 80 individual ants. RNA samples were sequenced using a dual-RNA sequencing approach ^47^, and the resulting reads were quality filtered and mapped to either the *A. echinatior* genome ^48^ (∼98% of reads, Supplementary Table 2) or their corresponding *Pseudonocardia* reference genomes^31^ (∼1% of reads, Supplementary Table 2). Both *Pseudonocardia* species showed very similar patterns of gene expression *in vivo*, with genes involved in the production of secondary metabolites, including antibiotics (as classified by KEGG) being expressed at similar levels by both *Pseudonocardia* strains on the cuticle of *A. echinatior* ants (Supplementary Figure 7).

BGCs that are shared by the *P. octospinosus* and *P. echinatior* strains^31^ (Supplementary Table 3) displayed remarkably similar patterns of *in situ* expression on the propleural plates (Figure 4A). For both *Pseudonocardia* species, the most highly expressed BGCs encoded proteins responsible for the synthesis of the compound ectoine and a putative carotenoid terpene pigment (cluster D and F, Figure 4A). Such compounds are known to provide protection against abiotic stressors such as desiccation and high concentrations of free radicals which are often associated with biofilms^49, 50, 51^. Also expressed on the propleural plates is a shared BGC encoding a putative bacteriocin (Cluster E, Figure 4A), which belongs to a family of ribosomally-synthesized post-translationally modified peptide (RiPP) antibiotics produced by many species of bacteria^52, 53, 54^; these are known to prevent the formation of biofilms by other microbial species^52, 55^. In contrast, a shared Type 1 PKS gene cluster, encoding nystatin-like antifungal compounds^31, 37^, had very low expression in the *P. echinatior* strain and was not expressed at all in the *P. octospinosus* strain (cluster C, Figure 4A). This suggests that additional cues, such as direct exposure to *E. weberi* may be required to activate this BGC, since both *Pseudonocardia* strains produce inhibitory antifungals when confronted with *E. weberi in vitro* (Supplementary Figure 8). A *Pseudonocardia* strain isolated from *Acromyrmex* ants has also previously been shown to produce nystatin-like compounds *in vitro*^37^. The most highly expressed BGCs unique to *P. echinatior* (cluster P, Figure 4C) or *P. octospinosus* (cluster J, Figure 4B) are both predicted to encode bacteriocins.

**Figure 4.**
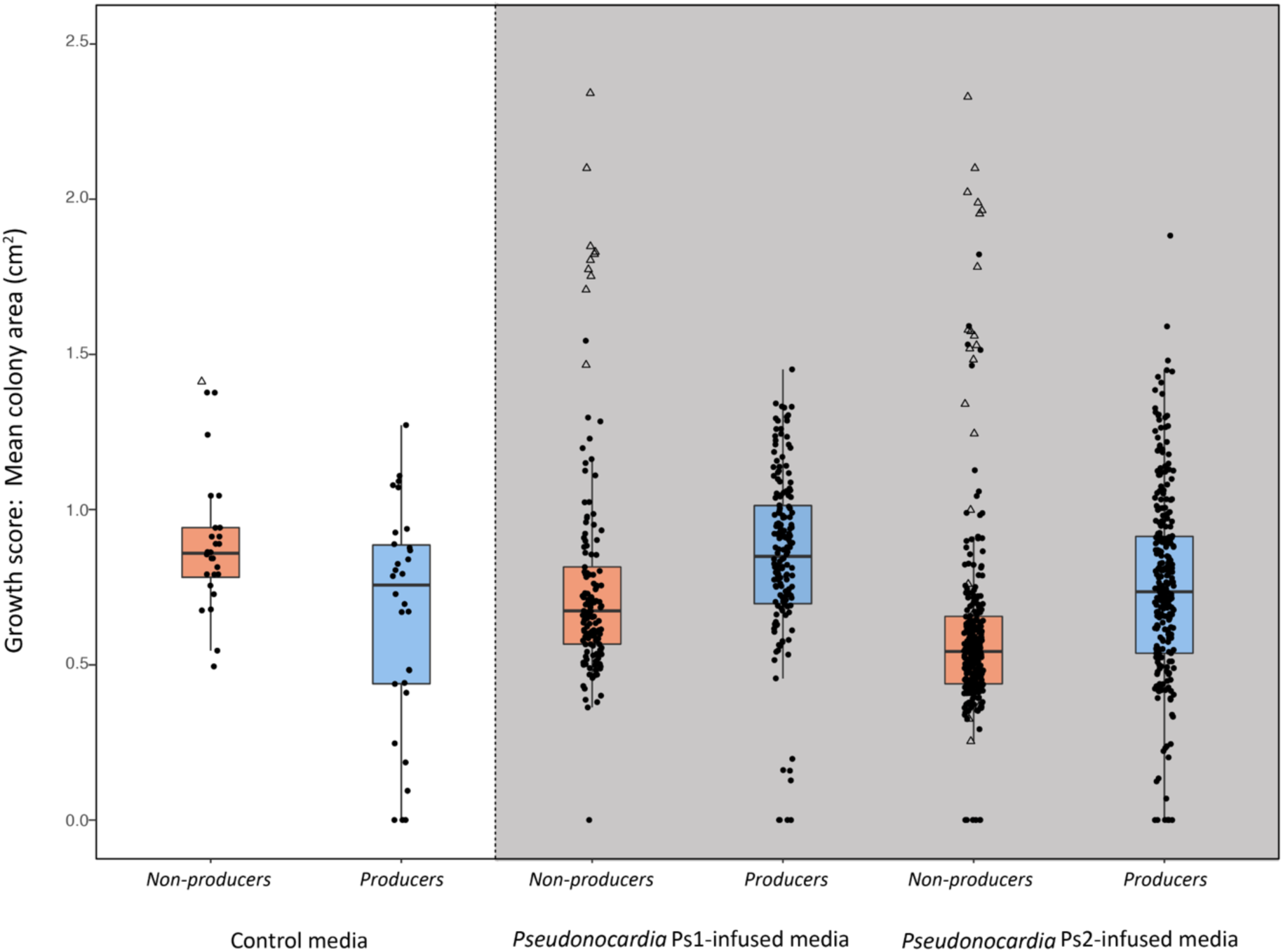
Growth-rate experiments. Bacterial colony sizes after 5 days at 30 °C. Boxplots indicate medians ± one quartile. The white section shows growth rates on control media, and the grey section shows growth rates on the *Pseudonocardia*-infused media. Red boxes represent non-antibiotic-producer strains, and blue boxes represent antibiotic-producing *Streptomyces* strains. A linear mixed-effects model ^56^, including 17 *Pseudonocardia* strains and 20 inoculated bacterial species as random intercepts, was used to test for interaction and main effects of growth media (Control vs. Ps1.infused vs. Ps2.infused) and antibiotic production (Non.producers vs. Producers): *lme4::lmer*(Growth.score ∼ Growth.media*Antibiotic.production+(1|Ps.strain/Plate)+(1|In.strain). One non-producer strain (*Staphylococcus epidermidis*) grew more rapidly than all other strains (open triangle points), demonstrating the need to control for correlated residuals.

Taken together, the results of the RNA-SIP and dual-RNA sequencing experiments are consistent with both previous empirical research^18, 32^ and the screening hypothesis^11, 12^: the ant host provides public resources to its cuticular microbiome via glandular secretions^40^ for which additional ectosymbionts compete once workers become exposed. This is always after the native *Pseudonocardia* mutualists have established and gained dominance, which invariably creates a demanding cuticular environment for any additional strain to invade.

### *Pseudonocardia* antibacterials create a demanding environment for non-antibiotic-producing bacteria

Next we compared the growth rates of antibiotic-producing *Streptomyces* strains and non-antibiotic-producing bacteria, which were all isolated from soil or fungus-growing ant nests (Supplementary Table 1), on antibiotic-infused and control media. The antibiotic-infused media were created by growing lawns of 17 *Pseudonocardia* isolates (Table 1) on SFM agar, while control media were inoculated with 20% glycerol. After a six-week incubation period, the agar medium was flipped to reveal a surface for colonisation. The non-producer strains grew more quickly on the undemanding control media while the antibiotic-producers grew more quickly on the demanding *Pseudonocardia*-infused media, producing a highly significant statistical interaction effect (n = 975, χ^2^ = 45.86, df = 2, p < 0.0001; Fig. 4). There was also a significant main effect of *Pseudonocardia* genotype, with both non-producers and producers exhibiting a lower growth rate on *P. echinatior* (Ps2)-infused media than on *P. octospinosus* (Ps1)-infused media (linear mixed-effects model, n = 915, χ^2^ = 24.55, df = 1, p < 0.0001, control-media data omitted for this analysis). This outcome is consistent with the observation by Andersen *et al*. ^18^ that *Acromyrmex* colonies hosting Ps2-dominated cuticular microbiomes were less prone to secondary invasion by other bacteria.

To test the hypothesis that producer strains are generally resistant to antibiotics, which would confer competitive superiority in an antibiotic-infused host environment^11^, we grew ten producer strains (all *Streptomyces* spp.) and ten non-producer strains (Supplementary Table 1) in the presence of eight different antibiotics (Supplementary Table 4), representing a range of chemical classes and modes of action. After seven days, Lowest Effective Concentration (LEC, lowest concentration with inhibitory effect) and Minimum Inhibitory Concentration (MIC, lowest concentration with no growth) scores were assigned on a Likert scale of 1–6, where a score of 1 was no resistance and a score of 6 was resistance above the concentrations tested^57^. Antibiotic-producer strains exhibited greater levels of resistance, measured by both LEC (Wilcoxon two-sided test, W = 94.5, *p* = 0.0017) and MIC scores (W = 80, *p* = 0.0253); p-values corrected for two tests (Fig. 5).

**Figure 5.**
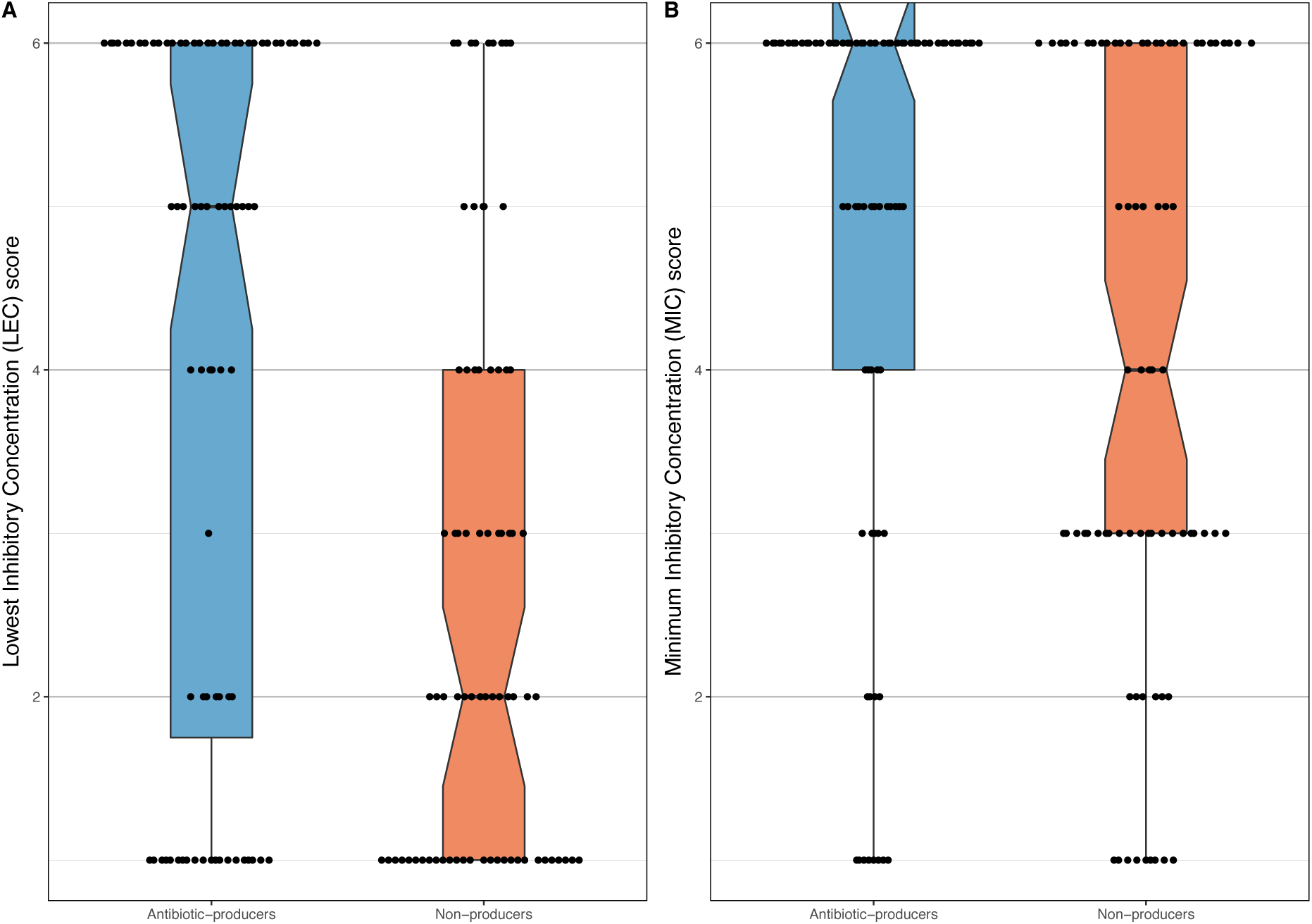
Antibiotic resistance profiles (**A** LEC and **B** MIC) for *producer*s and *non-producer* strains (Supplementary Table 1), based upon each strain’s mean growth score across the eight tested antibiotics (details in S3). Boxplots indicate medians (notches) ± one quartile. ‘Rabbit ears’ in **B** indicate that the medians are also the highest values.

We also performed growth rate experiments and measured antibiotic resistance profiles with resident non-producer strains that were directly isolated from cuticular microbiomes. These strains had significantly slower growth rates overall, even on control media without antibiotics, suggesting that they may only be transient environmental contaminants (Supplementary Figure 9). Resident non-producer strains also demonstrated high levels of resistance (Supplementary Figure 10) as expected, given that they were isolated from ant cuticles.

### *Pseudonocardia* antibacterials allow *Streptomyces* to competitively exclude non-antibiotic-producing bacteria

Finally, to test whether producer strains have a competitive advantage in the demanding environment created by *Pseudonocardia*, we pairwise-competed two of the *Streptomyces* producer strains, named S2 and S8 (Supplementary Table 1), against each of 10 environmental non-producer strains on normal and on antibiotic*-*infused media. In the latter case, *Pseudonocardia* was again grown on agar plates before turning the agar over and coinoculating producer and non-producer test strains. On normal growth media, *Streptomyces* were more likely to lose to non-producers, but when grown on *Pseudonocardia*-infused media, *Streptomyces* were more likely to win (S8 on Ps1-infused media: n = 129, χ^2^ = 103.6, df = 1, p < 0.0001; S2 on Ps2-infused media: general linear mixed-effects model, n = 94, χ^2^ = 87.9, df = 1, p < 0.0001; Figure 6).

**Figure 6.**
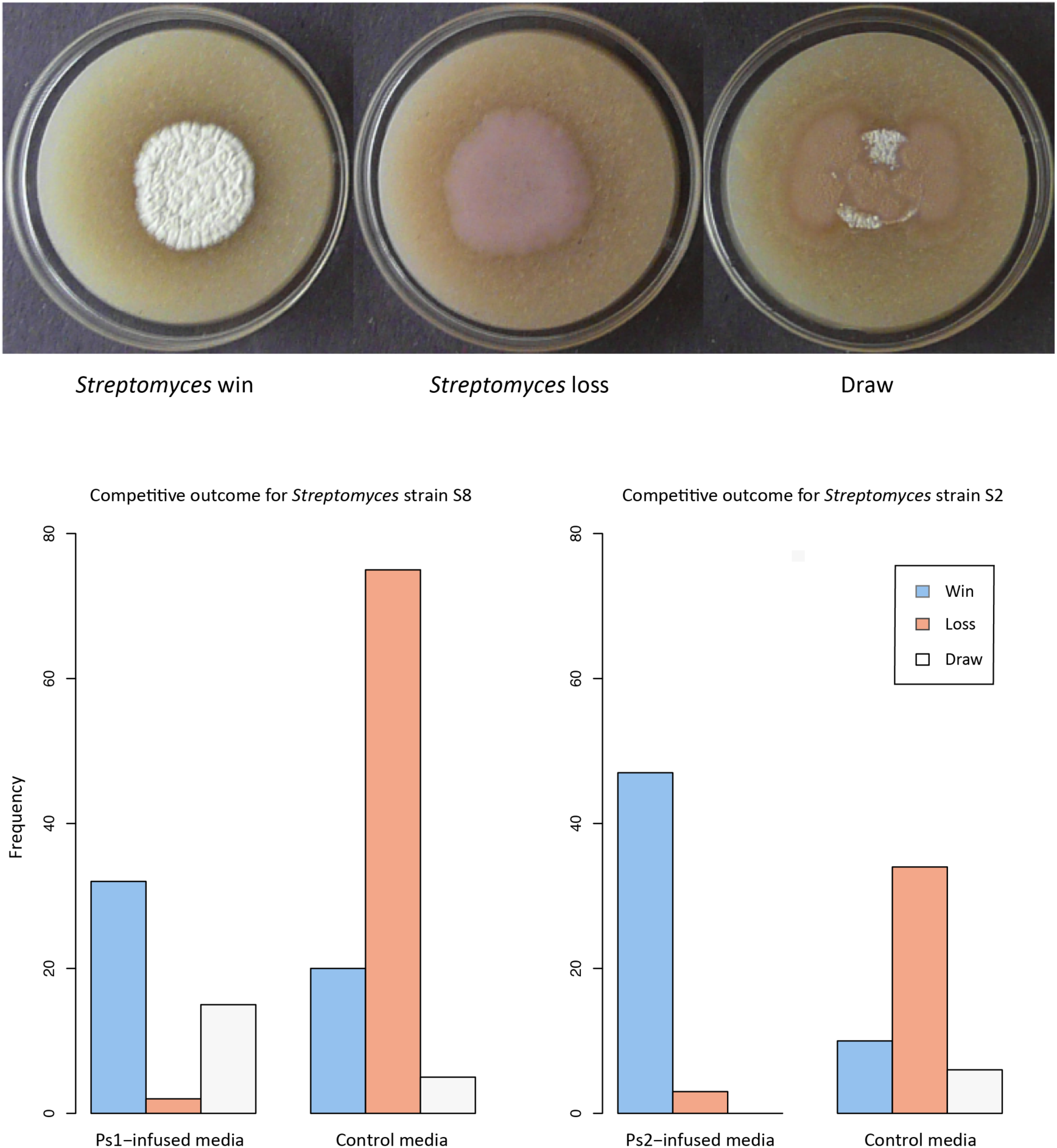
Pairwise competition experiment, scoring the frequency of producer wins. (**A**) Representative images of agar plates at five days post-inoculation showing examples of the three competitive outcomes: Win (producer S2 vs. non-producer St3 on Ps2 media), Loss (producer S8 vs. non-producer St3 on control media), and Draw (producer S2 vs. non-producer St3 on control media) (strain details in Supplementary Table 1). (**B**) Barcharts of competitive outcomes for the two *Streptomyces* producer strains (S8 and S2; Supplementary Table 1). Each *Streptomyces* strain was individually competed against ten different non-antibiotic-producer strains. *Streptomyces* is more likely to win on *Pseudonocardia-*infused media. A general linear mixed-effects model, with ten non-antibiotic-producer strains as a random intercept, was used to test for effect of medium (Control vs. Ps-infused) on competitive outcome (Win vs. Loss) *lme4::glmer*(outcome ∼ medium + (1 | non.producer.strain), family = binomial).

## Discussion

We tested screening theory using the external (cuticular) microbiome of the leafcutter ant *Acromyrmex echinatior* as an experimental model. We used RNA-SIP to show for the first time that an animal host is directly feeding its microbiome by secreting a public resource that all surface-resident bacteria can use (Figure 2), and we confirm that the public resource is indeed used by a number of environmentally acquired bacterial species. We further demonstrated, in two separate captive colonies, that both *P. octospinosus* and *P. echinatior* strains express antibacterial biosynthetic gene clusters (BGCs) on the ant cuticle (Figure 3). We next showed that the two species of actinobacteria have broad-spectrum antibacterial activity against environmental isolates *in vitro* but, importantly, have a weaker effect on *Streptomyces* species (Figure 4), consistent with these *Streptomyces* species being resistant to a range of antibiotics, which is typical for this genus (Figure 5). Finally, we used *in vitro* competition experiments to demonstrate that the *Streptomyces* species can competitively exclude faster-growing bacteria that do not make antibiotics, but only when the competing species are grown on media infused with *Pseudonocardia* metabolites (Figure 6).

**Figure 3.**
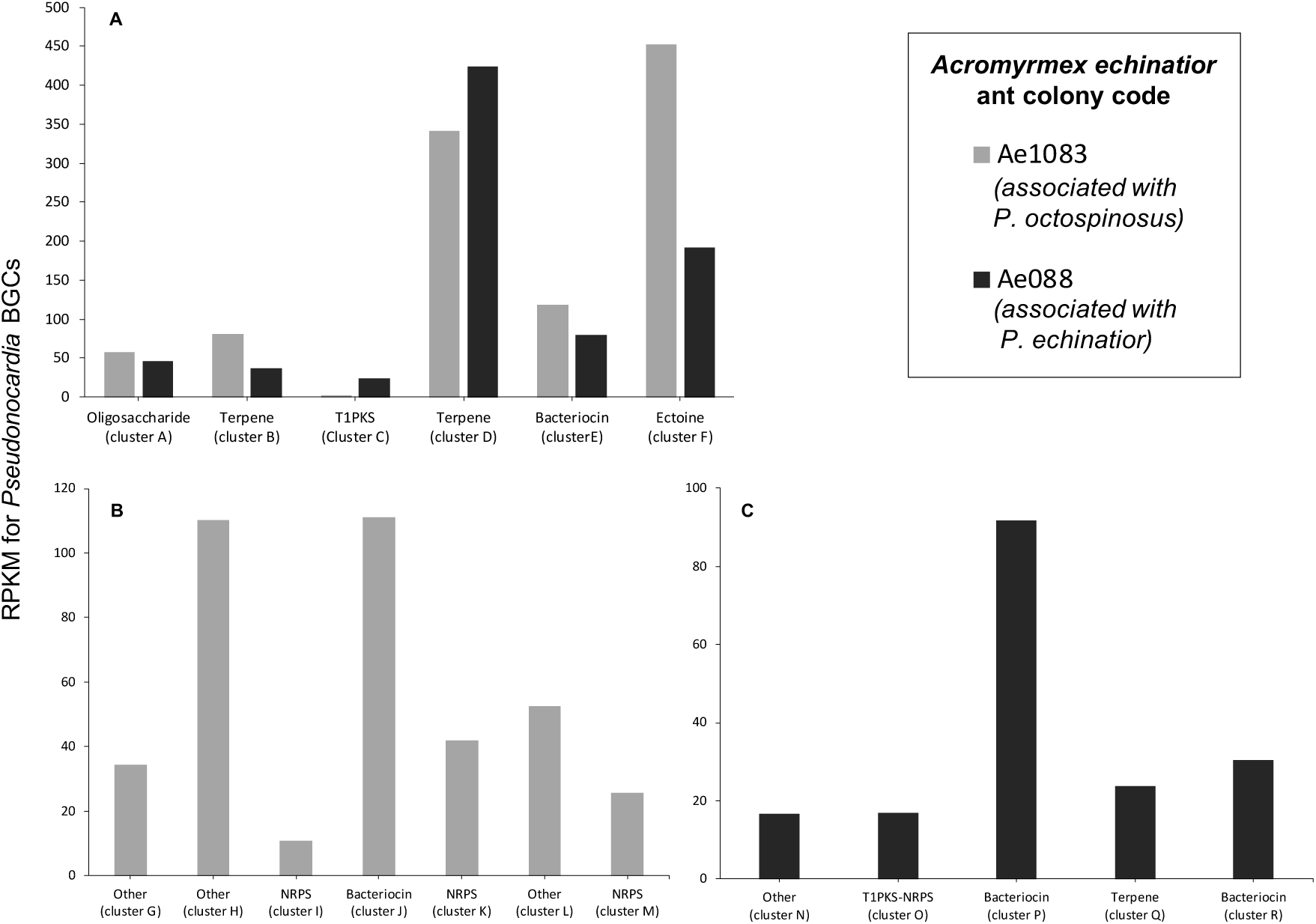
Expression (in reads per kilobase of transcript per million mapped reads, RPKM) of (**A**) biosynthetic gene clusters that are shared between the two *Pseudonocardia* species, (**B**) biosynthetic gene clusters that are unique to *P. octospinosus* (colony Ae1083) and (**C**) biosynthetic gene clusters that are unique to *P. echinatior* (colony Ae088). Biosynthetic gene cluster codes relate to Table S3. Results from each colony were derived from a pool of 80 ants.

Taken together, these results are consistent with the hypothesis that competition-based screening is a plausible mechanism for the environmental acquisition and assembly of a multi-species defensive microbiome, provided that the vertically transmitted *Pseudonocardia* symbionts become established first to create a demanding environment. This priority establishment is invariably the case because callow large workers are inoculated by their nestmates within 24 hours of emerging from their pupae^34^, meaning that *Pseudonocardia* is never competitively excluded from the microbiome, as predicted by Scheuring & Yu ^11^ and empirically shown by Andersen *et al.* ^18^. The new results reported here confirm the tractability of the cuticular leafcutter-ant microbiome, which is accessible to experimentation and for which the adaptive benefit to the ant hosts is clearly defined and explicitly testable: defence against specialized *Escovopsis* pathogens^25, 37, 58, 59^. Our present results remain consistent with *Streptomyces* strains being superior contenders for secondary acquisition, in the context of, but without being a threat to, the vertical transmission across generations of the native symbionts by dispersing gynes that found new colonies after being inseminated.

Our results are of broad significance because they provide mechanistic evidence for the combined applicability of the evolutionary-ecosystem-on-a-leash concept^7^ and screening theory specifying the ecological dynamics of tilting the competitive landscape in favour of other antibiotic-producing bacteria^11^. The evolutionary perspective suggests that, depending on their degree of openness, host-associated microbiomes may have both native core members that co-adapt with host environments, while the ecological perspective addresses the mechanism by which successfully establishing bacterial colonists are unlikely to be parasites or free-riders. Studies in other symbioses appear to support this dual evolutionary and ecological view^13, 15, 16^, both for an array of mutualistic symbioses with multicellular partners and for microbiomes more specifically. The combined evolutionary and ecological perspective of our present study is remarkably parallel to the priority effects documented for vertically transmitted (actinomycete) *Bifidobacterium* species in the milk glycobiomes that inoculate the gut microbiome of human neonates^10, 60^. Similar processes may also play out during the establishment of plant-root microbiomes^61, 62, 63^ and could be open to manipulation in efforts to improve crop yields^64^.

Future work on the *Acromyrmex* model system should aim at characterising the antibacterial molecules made by the symbiotic *Pseudonocardia* strains *in vitro* and *in vivo* and matching these compounds to the BGCs expressed on the ants. This will be challenging because ant-derived *Pseudonocardia* strains grow poorly on agar plates and not at all in liquid culture. Additionally, Imaging Mass Spectrometry has, as yet, not been possible on the ants themselves.

## Materials and Methods

### Ant colony collection and maintenance

Colonies of *A. echinatior* (Hymenoptera, Formicidae, Attini) were collected from the Gamboa area of the Soberania National Park, Panama, between 2001 and 2014. Colonies Ae1083 and Ae088 (Supplementary Table 1) were maintained under controlled temperature conditions (25°C) at UEA and fed a daily diet of bramble and laurel leaves. Additional colonies were maintained at the University of Copenhagen in rearing rooms at ca 25°C and 70% relative humidity, where they were fed with bramble leaves and occasional supplements of apple and dry rice.

### RNA SIP ^13^C feeding experiment

Six replicate groups of 22 mature worker ants with visible bacterial growth on their propleural plates were selected from colony Ae1083 (Supplementary Table 1) and placed into 9cm petri dishes containing a 2×2 cm square of cotton wool soaked in water. Following 24 hours of starvation, three replicate groups of 22 ants were supplied with 300 μl of a 20% ^13^C glucose solution (w/v, Sigma Aldrich) and the remaining three groups were supplied with 300 μl of a 20% ^12^C glucose solution (w/v, Sigma Aldrich) for 10 days. Glucose solutions were supplied to ants in microcentrifuge tube caps and were refreshed every three days. To confirm uptake of the ^13^C isotope by ants fed on the ^13^C labelled diet, a further 5 ants were fed on each type of glucose diet; these were submitted for Isotope Ratio Mass Spectrometry (IRMS) analysis (Supplementary Note 1), which enables the relative abundance of each stable isotope (^13^C and ^12^C) to be quantified in a sample. To confirm that the glucose water diet did not spread over the ant cuticle, a 20% glucose solution containing a non-toxic fluorescent green drain tracing dye (5 mg ml^-1^, Hydra) was fed to the ants. Ants were sampled just after taking a feed, and after 6 and 24 hours of being exposed to the dye, to trace the spread of the solution over time. The ants were imaged using fluorescent microscopy (Supplementary Figure 2, Supplementary Note 1).

### Cuticular dissection and RNA extraction

At the end of the 10-day feeding experiment, the propleural plates of the propleura were removed from the ventral exoskeleton using a dissection microscope and fine sterile tweezers. Propleural plates from each of the 22 ants in each group were placed together in lysis matrix E tubes (MP Biomedicals) on dry ice, before being snap frozen in liquid nitrogen. A modified version of the Qiagen RNeasy Micro Kit protocol was used for all RNA extractions (Supplementary Note 1). The quantity and purity of all RNA samples was checked using a nanodrop spectrophotometer and a Qubit™ RNA HS assay kit (Invitrogen™).

### Density gradient ultracentrifugation and fractionation

Density gradient ultracentrifugation was carried out to separate ^13^C labelled (“heavy”) from un-labelled (“light”) RNA within the same sample (Supplementary Note 1). Following centrifugation, samples were divided into 12 fractions using a peristaltic pump to gradually displace the gradient, according to an established protocol^43^. The refractive index of fractions was measured to confirm the formation of a linear buoyant density gradient (Supplementary Figure 3). RNA was precipitated from fractions by adding 1 volume of DEPC-treated sodium acetate (1M, pH 5.2), 1 μl (20 μg) glycogen (from mussels, Sigma Aldrich) and 2 volumes of ice cold 96% ethanol. Fractions were incubated over night at -20°C then centrifuged for 30 minutes at 4°C and 13,000 x g, before washing with 150 μl of ice cold 70% ethanol and centrifuging for a further 15 minutes. Pellets were then air-dried for 5 minutes and re-suspended in 15 μl of nuclease free water.

### Quantifying 16S rRNA gene copy number across RNA SIP fractions

The RNA in each fraction was converted to cDNA by following the manufacturer’s instructions for Superscript II (Invitrogen) with random hexamer primers (Invitrogen). 16S rRNA gene copy number was then quantified across cDNA fractions using qPCR (Supplementary Note 1).

### Sequencing and analysis

PCR was used to amplify the 16S rRNA gene in each of the fractions that spanned the peaks in 16S rRNA gene copy number, identified via qPCR. This was done using the primers PRK341F and 518R (Supplementary Table 1). One unfractionated sample was also created for ants under each of the ^13^C or ^12^C dietary treatments, by pooling equal quantities of unfractionated cDNA from each of the 3 replicate groups and using this as template for PCR amplification. The resulting PCR products were purified using the Qiagen MinElute™ gel extraction kit and submitted for 16S rRNA gene amplicon sequencing using an Illumina MiSeq at MR DNA (Molecular Research LP), Shallowater, Texas, USA. Sequence data was then processed by MR DNA using their custom pipeline^65, 66^. As part of this pipeline, paired-end sequences were merged, barcodes were trimmed, and sequences of less than 150 bp and/or with ambiguous base calls were removed. The resulting sequences were denoised, and OTUs were assigned by clustering at 97% similarity. Chimeras were removed, and OTUs were taxonomically assigned using BLASTn against a curated database from GreenGenes, RDPII, and NCBI^67^. Plastid-like sequences were removed from the analysis. Upon receipt of the 16S rRNA gene sequencing data from MR DNA, OTU assignments were verified using QIIME2 and BLASTn, and statistical analysis was carried out using R 3.2.3^68^. OTUs assigned as *Pseudonocardia* were blasted against the 16S rRNA gene sequences for *P. echinatior* and *P. octospinosus* ^31^ to confirm the relative abundance of each of these vertically-transmitted strains in the samples. All 16S rRNA gene amplicon sequencing data from this experiment has been deposited in the European Nucleotide Archive (ENA) public database under the study accession number PRJEB32900.

### Detecting the expression of *Pseudonocardia* BGCs *in situ*

#### RNA extraction and sequencing from ant propleural plate samples

The propleural plates of *Acromyrmex echinatior* ants were dissected (as described above) from individual mature worker ants that had a visible growth of *Pseudonocardia* bacteria on their cuticle. A pool of 80 ant cuticles were sampled from each of the colonies Ae1083 and Ae088, respectively (Supplementary Table 1), after which RNA was extracted as described above. The quantity, purity and integrity of all RNA samples was checked using a nanodrop spectrophotometer and Qubit™ RNA HS assay kit (Invitrogen™), as well an Experion™ bioanalyser with a prokaryotic RNA standard sensitivity analysis kit (Bio-Rad, California, USA). One μg of RNA from each of the propleural plate samples was sent to Vertis Biotechnologie AG (Freising-Weihenstephan, Germany) where samples were processed and sequenced using a Dual RNA sequencing approach as described in ^47^. Single-end sequencing (75 bp) was performed using an Illumina NextSeq500 platform. All sequencing reads have been deposited in the ENA public database under the study accession number PRJEB32903. Upon receipt, the resulting RNA reads from each sample were quality filtered and aligned to both the *A. echinatior* reference genome^48^ and a representative genome for either *P. octospinosus* (from Ae707), or *P. echinatior* (from Ae706) (Supplementary Table 1; Supplementary Note 1). Reads that mapped to the *Pseudonocardia* genomes were then mapped back to the ant genome (and vice versa) to check that reads did not cross-map between the two genomes. Following alignment, uniquely mapped reads were counted and read counts per annotated coding sequence were converted to reads per kilobase of transcript per million mapped reads (RPKM) (Supplementary Note 1).

### Expression analysis

In order to investigate the expression levels of different functional groups of genes, protein sequences of every annotated gene in each *Pseudonocardia* genome^31^ (Supplementary Table 1) were extracted and uploaded to BlastKOALA^69^. Assigned K numbers were classified into five main KEGG pathway categories (and their associated sub-categories) using the KEGG Pathway Mapper tool. Each gene, with its associated K number and category assignments, was then matched to its RPKM value from the RNA sequencing dataset so that the expression levels of different KO categories could be established. To investigate the expression of biosynthetic gene clusters (BGCs) by *Pseudonocardia* on the ant cuticle, reference genome sequences for *P. octospinosus* and *P. echinatior*^31^ (Supplementary Table 1) were uploaded to antiSMASH version 4.0, which predicts the presence and genomic location of BGCs based on sequence homology to known clusters^70^. RPKM values were then generated for each predicted BGC, based on the length of the predicted cluster and read counts for genes situated within it.

### Escovopsis bioassays

The *Pseudonocardia* strains PS1083 and PS088 (Supplementary Table 1) were isolated from the propleural plates of individual large *Acromyrmex echinatior* workers taken from colonies Ae1083 and Ae088 (colonies used in RNA-seq experiments, see Supplementary Table 1), respectively. Each of the strains were bioassayed against the specialized fungal pathogen *Escovopsis weberi* strain CBS 810.71, which was acquired from the Westerdijk Fungal Biodiversity Institute (Supplementary Table S1). Isolation of the *Pseudonocardia* strains and bioassays are described in detail in Supplementary Note 1.

### Collection and isolation of bacterial strains

Nineteen strains of *Pseudonocardia* (11 strains of *P. echinatior* and 8 of *Pseudonocardia octospinosus,* Supplementary Table 1) were isolated from the cuticles of individual *Acromyrmex echinatior* worker ants across 18 different colonies and genotypes, as described in Supplementary Note 1. Ten of these isolated *Pseudonocardia* strains were previously genome-sequenced by Holmes *et al.*^31^. The 10 environmental antibiotic-producer strains (all in the genus *Streptomyces*) were taken from general collections in the Hutchings lab and are a mixture of isolates from either soil environments or from worker ants taken from captive colonies (Supplementary Table 1). The 10 environmental non-producer strains were obtained from the Hutchings lab (two strains) and from the ESKAPE suite (eight strains with varying origins (human skin, soil, etc.) used to test antibiotic resistance or efficacy in clinical/research settings (Supplementary Table 1).

### Individual growth rate and pairwise competition experiments

Antibiotic-infused media was created by growing lawns of 17 *Pseudonocardia* isolates (Supplementary Table 1) on SFM agar, whilst control SFM media was inoculated with 20% glycerol. After a six-week incubation period, the agar medium was flipped to reveal a new surface for colonisation. All *producer* and *non-producer* strains (Supplementary Table 1) were then inoculated onto the plates to compare their growth rates on each type of media (Supplementary Note 1). After 5 days of incubation at 30 °C, photographs were taken and growth area was measured. (Supplementary Note 1). Antibiotic-producer strains were also grown in direct competition against *non-producers,* both on normal control medium and on medium infused with *Pseudonocardia* secondary metabolites. Plates were incubated at 30 °C for 5 days before imaging. Images were then scored with respect to the *producer* as: win, draw or loss (Supplementary Note 1).

### Antibiotic resistance assays

To test whether producer and non-producer strains differed in their resistance to antibiotics, each strain (Supplementary Table 1) was grown in the presence of eight different antibiotics, representing a range of chemical classes and modes of action (Supplementary Table 4). Producers and non-producers were inoculated onto plates and incubated at 30 °C for 7 days, then photographed and scored on a Likert scale of 1–6. (Supplementary note 1).

### Statistical analyses

R scripts and data used for statistical analyses of Figures 4, 5, 6, and S9 are available at <u>github.com/dougwyu/Worsley_et_al_screening_test_R_code</u>

## Acknowledgements

This work was supported by a NERC PhD studentship to SFW (NERC Doctoral Training Programme grant NE/L002582/1), NERC grants NE/J01074X/1 to MIH and DWY, NE/M015033/1 and NE/M014657/1 to MIH, JCM, DWY, and BW, European Research Council Advanced grant 323085 to JJB, and Marie Curie Individual European Fellowship grant 627949 to TI. DWY was also supported by the Key Research Program of Frontier Sciences, Chinese Academy of Sciences (QYZDY-SSW-SMC024) and the Strategic Priority Research Program, Chinese Academy of Sciences (XDA20050202, XDB31000000). We thank the Smithsonian Tropical Research Institute for logistical help and facilities for all work in Gamboa, Panama, and the Autoridad Nacional del Ambiente y el Mar for permission to sample and export ants. We thank Simon Moxon for advice on RNA sequencing analysis and Paul Disdle for his assistentance with IRMS analysis. The authors declare no conflicts of interest.

## Supporting Information

**Supplementary Note 1:** Additional information regarding methods used for IRMS analysis, RNA extractions and the analysis of RNA sequencing, in addition to detailed methods for the growth-rate and pairwise competition experiments.

---

**Supplementary Figure 1.**
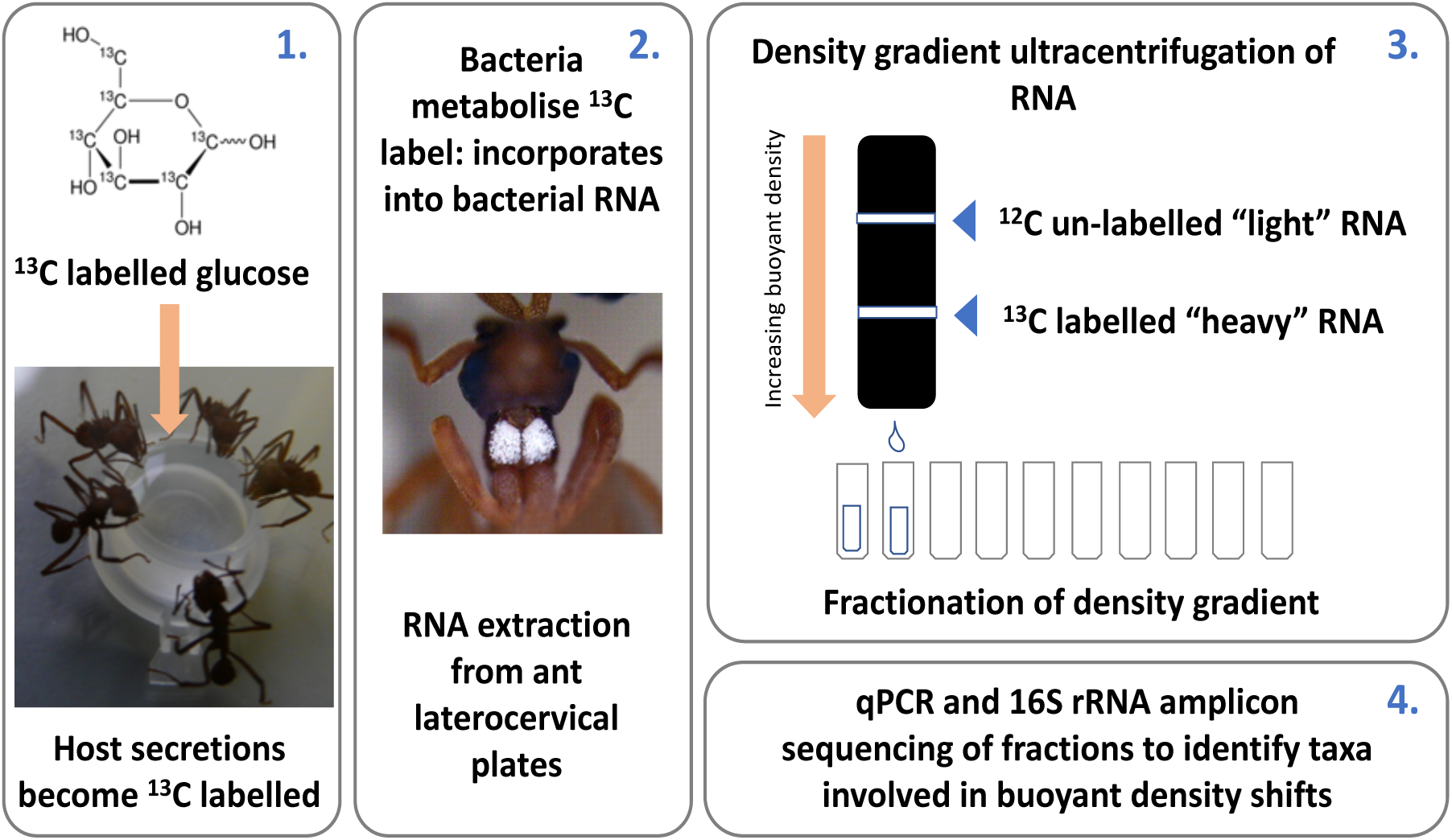
Overview of methodology used for RNA stable isotope probing of the propleural plate microbiome of *Acromyrmex echinatior* leafcutter ants.

**Supplementary Figure 2.**
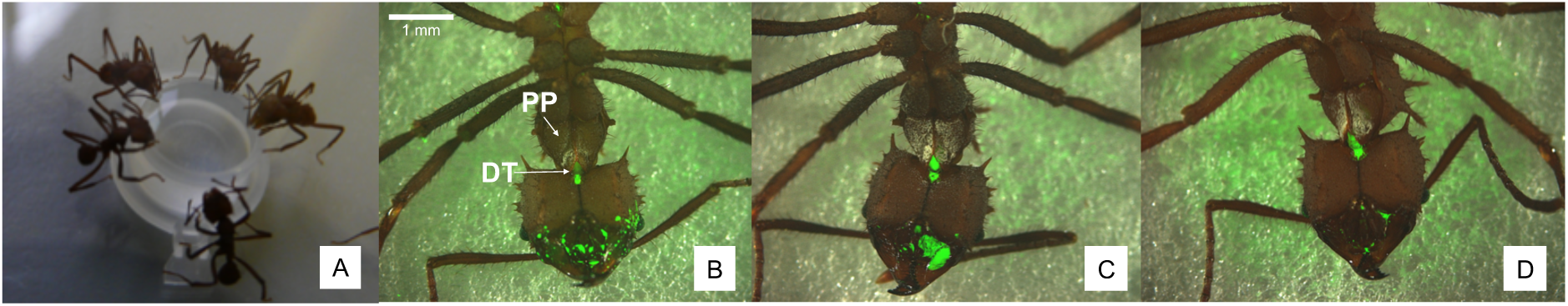
The extent to which a 20% (w/v) glucose water diet containing green fluorescent dye (A) was distributed over the ant body directly after taking a feed (B), 6 hours after feeding (C) and 24 hours after feeding (D). DT = fluorescence shining through from the ant digestive tract, PP = propleural plates with growth of actinobacteria.

**Supplementary Figure 3.**
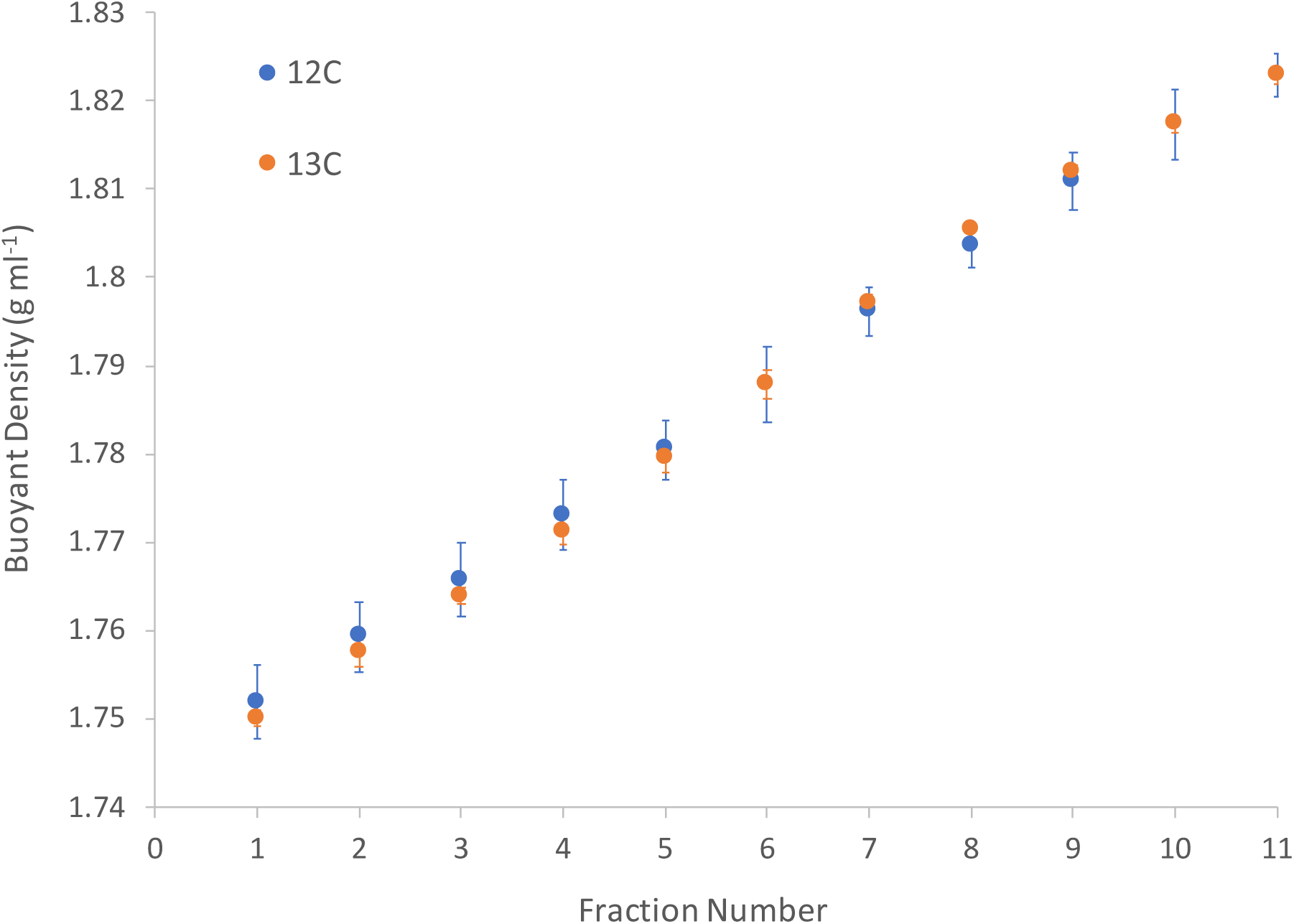
The relationship between RNA-SIP fraction number and buoyant density (in g ml^-1^). Fractions were generated from density gradients containing RNA isolated from the propleural plates of *A. echinatior* ants fed on either a ^12^C (blue) or a ^13^C (orange) glucose water diet. Points represent averages (three samples each of 22 ants per dietary treatment) ± standard error.

**Supplementary Figure 4.**
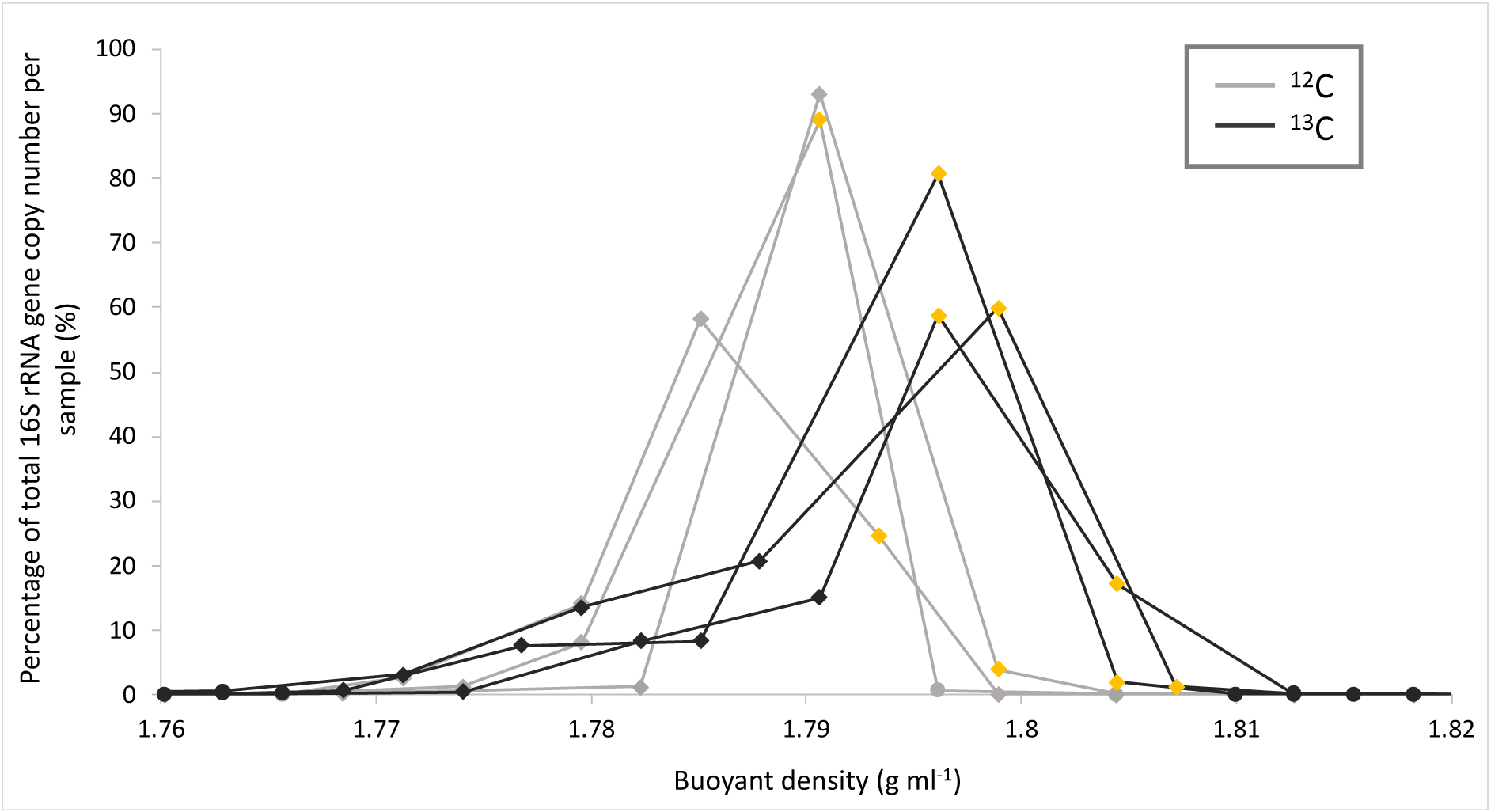
16S rRNA gene copy number across different fractions of buoyant density gradients, as determined via qPCR. Gene copy number in each fraction is displayed as a percentage of total copy number in each sample. There were three replicate samples of 22 ants fed a ^12^C (light grey) or ^13^C (dark grey) glucose diet. Diamond symbols represent fractions that were sent for 16S rRNA gene amplicon sequencing and yellow diamonds represent those designated as “heavy” fractions under the different treatments.

**Supplementary Figure 5.**
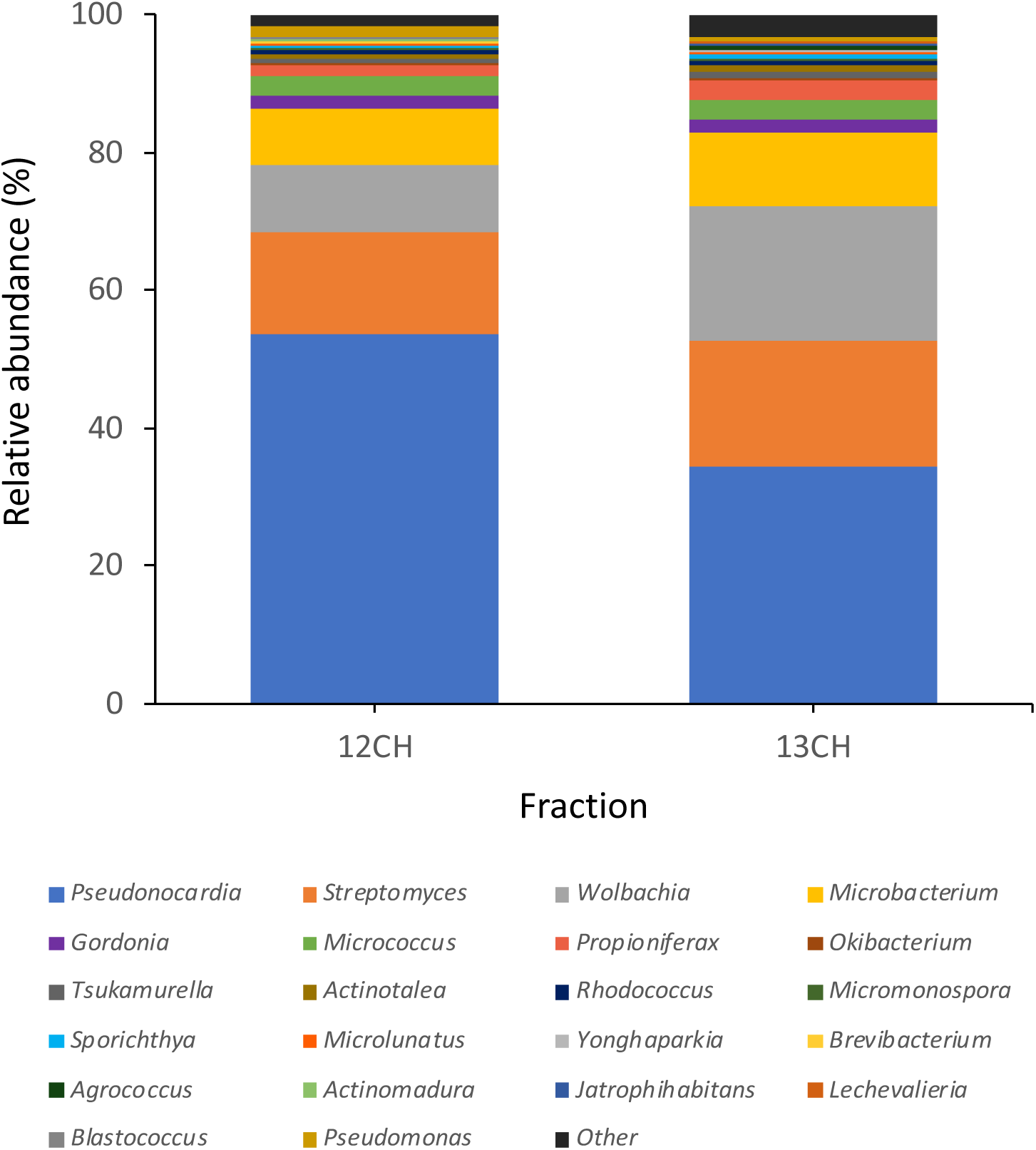
The relative abundances (%) of genera in the “heavy” fractions of buoyant density gradients for ^12^C and ^13^C samples, respectively. Heavy fractions were determined via qPCR, which identified peaks in 16S rRNA gene copy number in both sets of samples (see Supplementary Figure 4). Genera reported in Figure 1 (in unfractionated samples) were also recovered in fractionated samples; *Pseudonocardia* was the most prevalent genus, followed by *Streptomyces, Wolbachia* and *Microbacterium*. Additional actinomycete genera were recovered at lower abundances. Genera labelled as “other” were recovered in trace amounts.

**Supplementary Figure 6.**
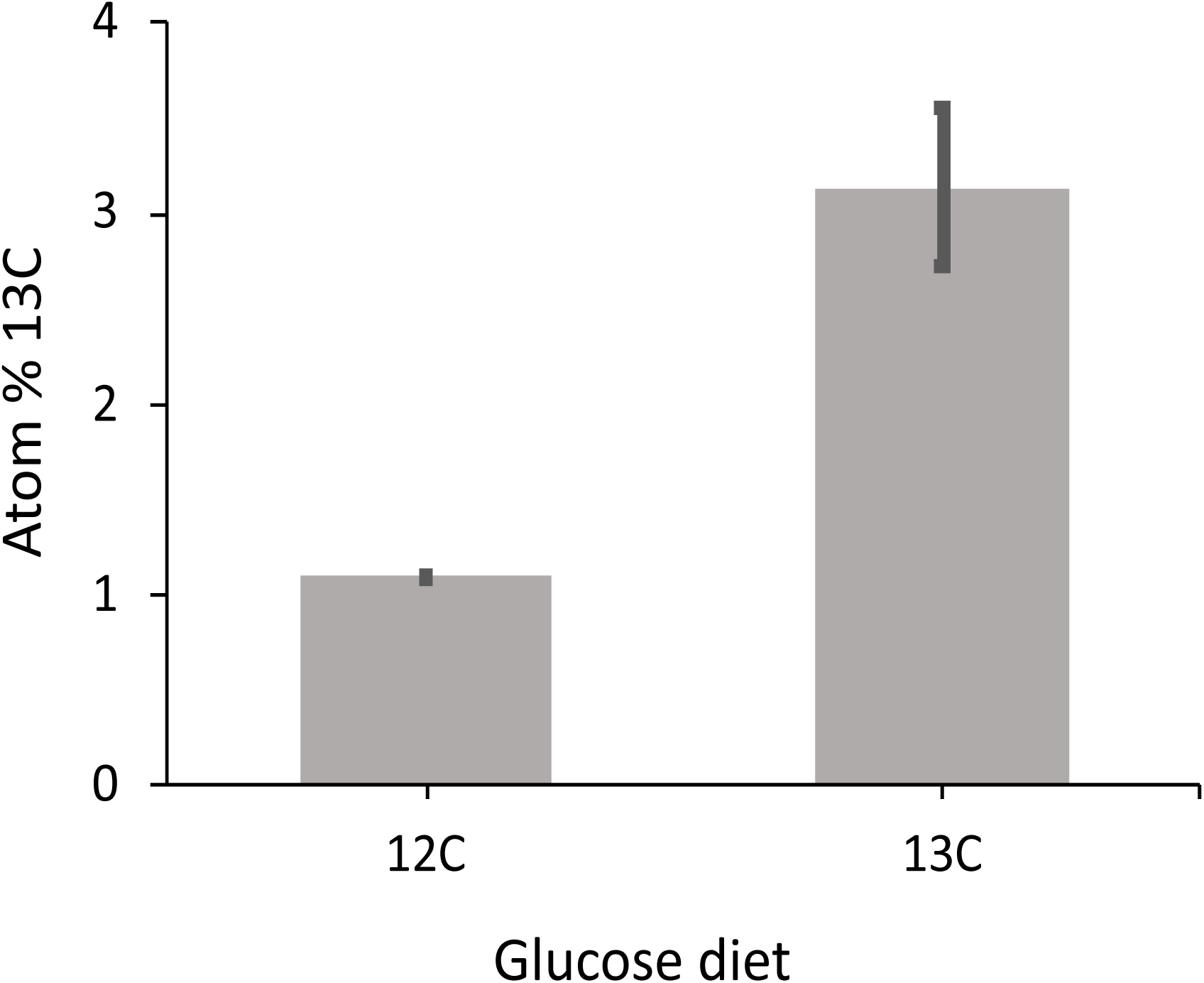
The atom percentage of ^13^C in ants fed either a ^12^C or ^13^C labeled 20% (w/v) glucose diet for 10 days, as determined by Isotope Ratio Mass Spectrometry (IRMS) analysis.

**Supplementary Figure 7.**
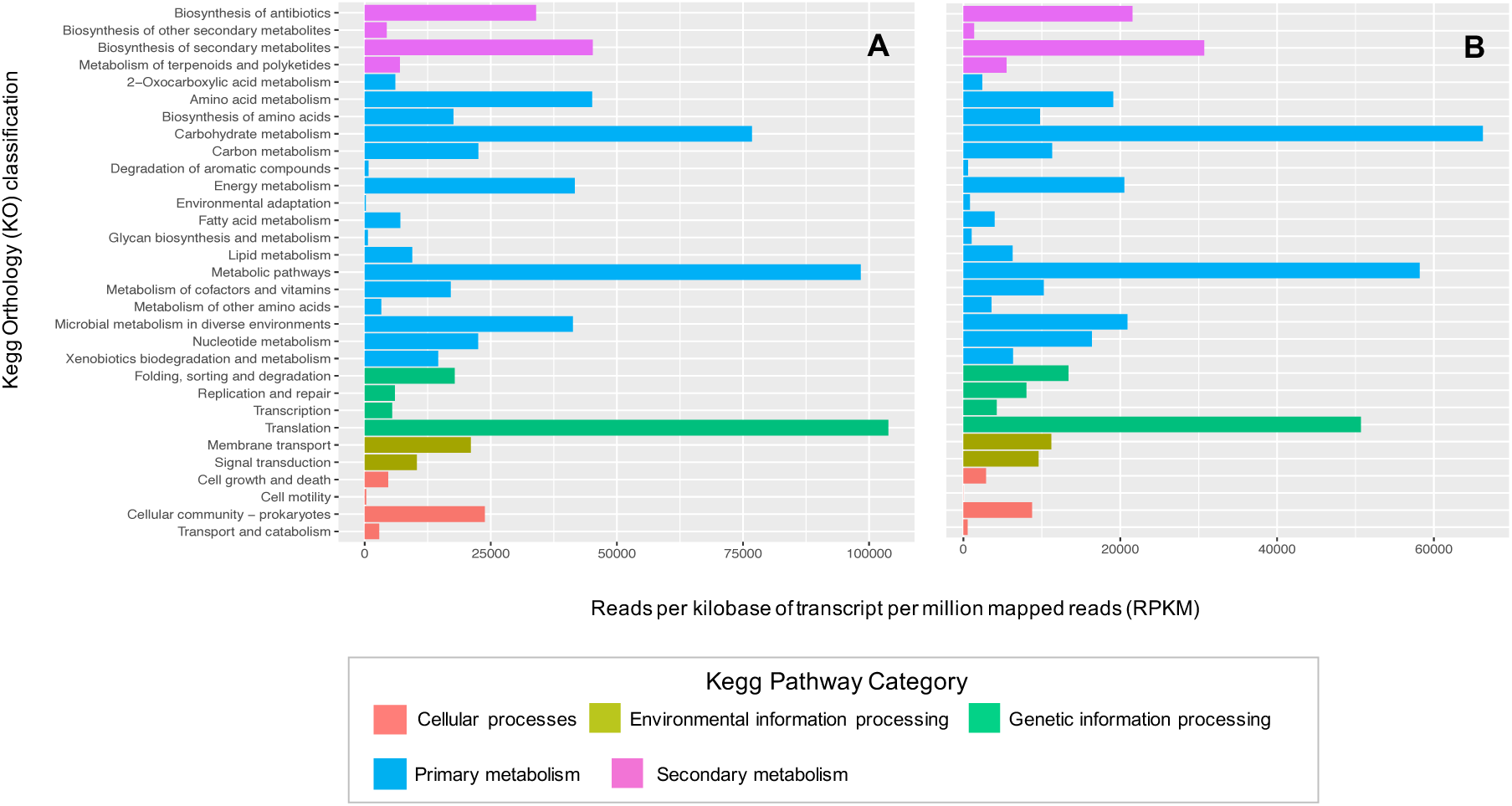
Expression levels (in reads per kilobase of transcript per million mapped reads, RPKM) of Kegg orthology pathway categories. (A) *Pseudonocardia octospinosus* (colony Ae088) and (B) *Pseudonocardia echinatior* (colony Ae1083) on the propleural plates of *Acromyrmex echinatior* ants. N= 1 sample of 80 pooled ants per colony.

**Supplementary Figure 8.**
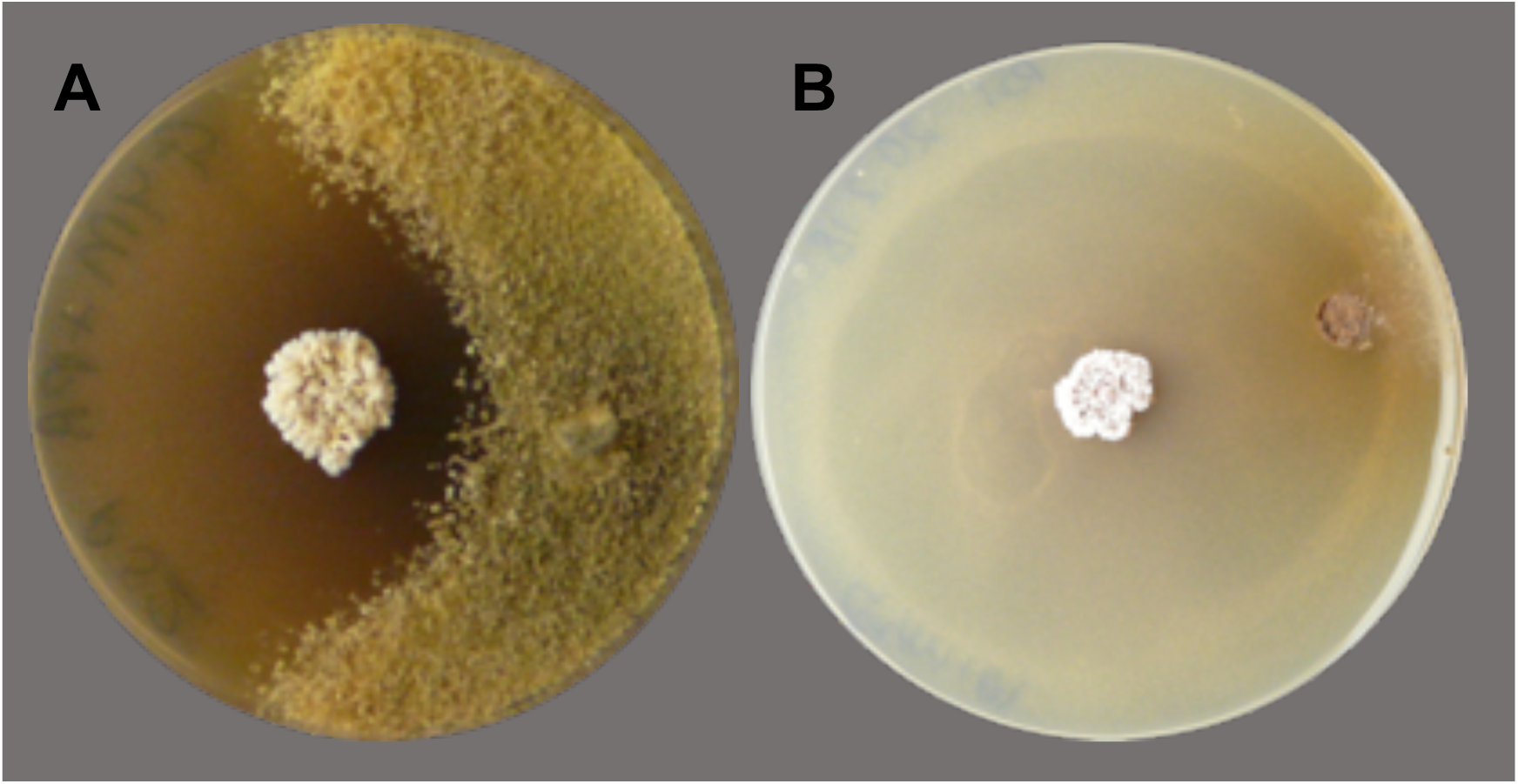
The bioactivity of *Pseudonocardia* isolates (**A)** *P. echinatior* PS088 and (**B)** *P. octospinosus* PS1083 against the specialized fungus-garden pathogen *Escovopsis weberi* (Supplementary Table 1).

**Supplementary Figure 9.**
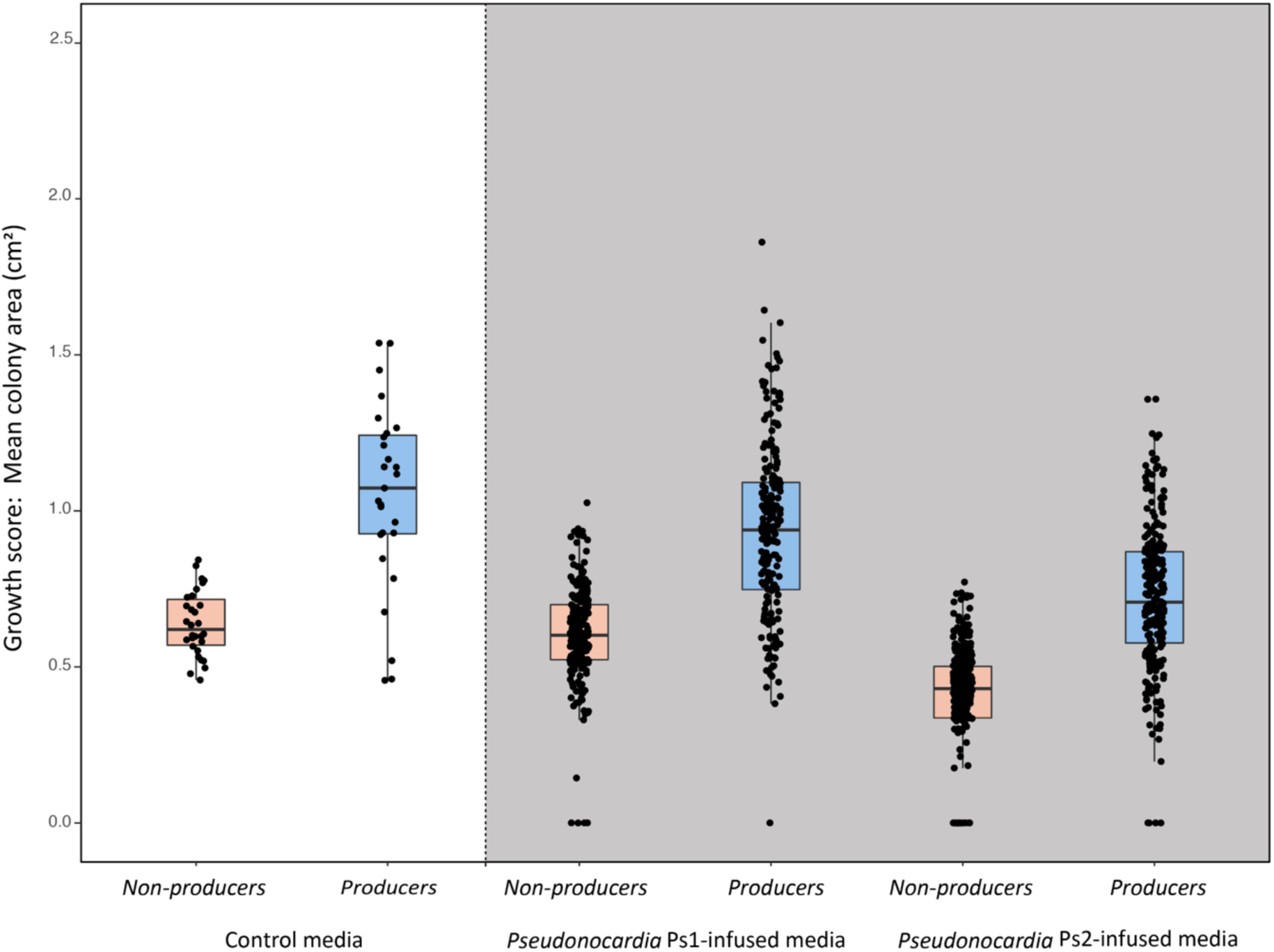
Individual growth-rate experiments of *Acromyrmex*-resident, *non-producer* strains, assessed as bacterial colony sizes after 5 days at 30 °C, with the boxplots indicating medians ± one quartile. The white section shows growth rates on control media, and the grey section shows growth rates on the *Pseudonocardia*-infused media. Red boxes represent *non-producer* strains, and blue boxes represent *producer* strains. For analysis, a linear mixed-effects model, including *Pseudonocardia* strain (n=17) and inoculated bacterial species (n=20) as random factors, was used to test for interactions and main effects of growth media (Control vs. Ps1-infused vs. Ps2-infused) and antibiotic production (Non producers vs. *Streptomyces*). There was no significant interaction effect (χ^2^ = 2.64, df = 2, p = 0.27), but both main effects were highly significant. The resident non-producers isolated from cuticular microbiomes had significantly slower growth rates on all media (χ^2^ = 20.96, df = 1, p < 0.0001) including the control media without antibiotics. This suggests that they are unable to outcompete producer strains on the cuticle of *Acromyrmex* ants and raises the question why these non-producer species can persist at all. Bacterial growth was also generally slower on Ps2-infused media than on Ps1-infused media (χ^2^ = 21.43, df = 1, p < 0.000 when analyzed in a balanced design without the control-media.

**Supplementary Figure 10.**
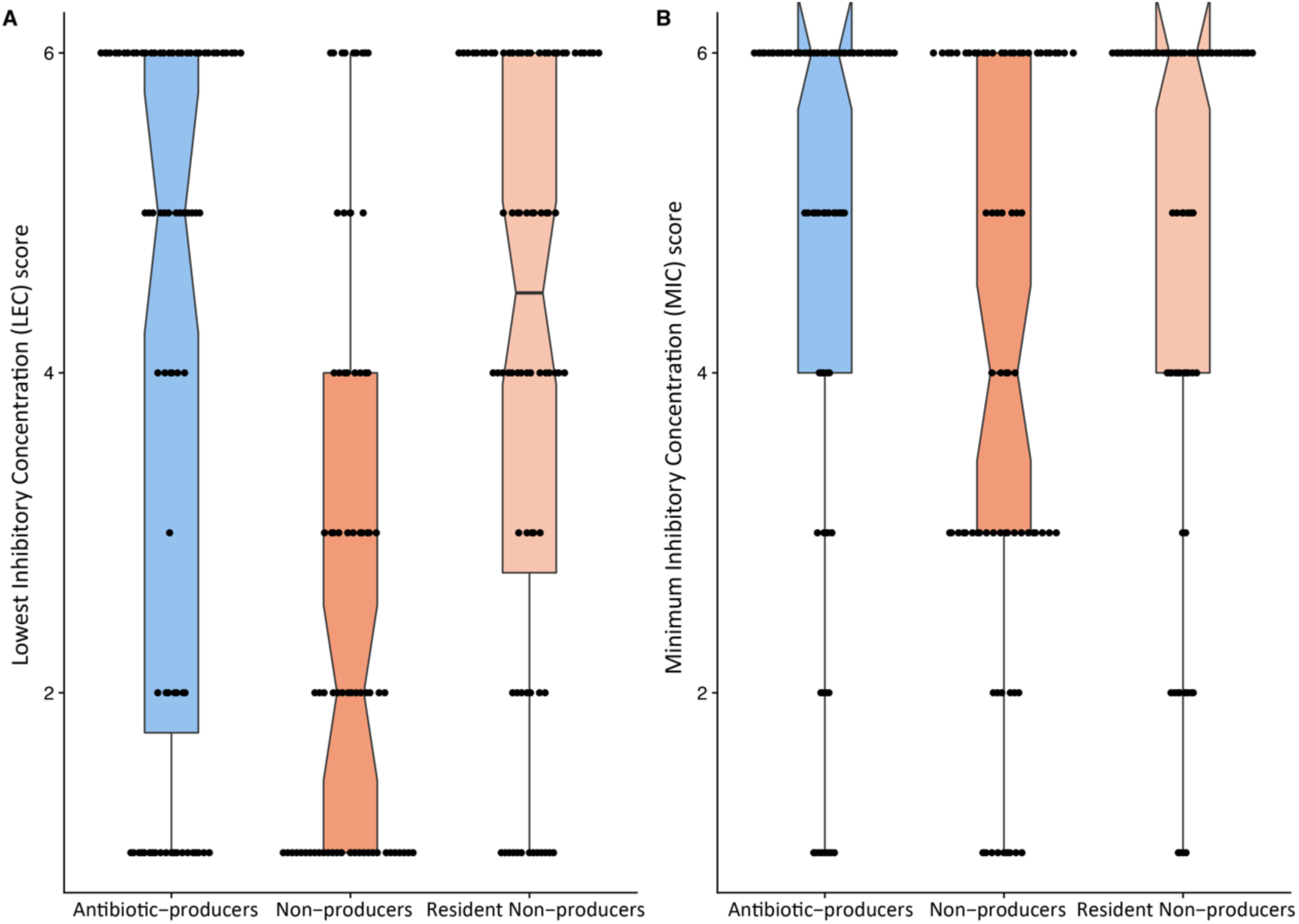
Antibiotic resistance profiles for *producer*, *non-producer*, and *resident non-producer* strains (Supplementary Table 1). Boxplots indicate medians (notches) ± one quartile. For analysis, we calculated each strain’s mean growth score across the eight tested antibiotics (reducing from n = 155 to n = 20; details in S3). *Producers* showed higher levels of resistance than did *non-producers* for both measurements: Wilcoxon two-sided test (wilcox.test), W = 94.5, p = 0.0017 for LEC (**A**) and W = 80, p = 0.0253 for MIC (**B**), after correction for multiple testing. *Producers* and *Resident Non-producers* showed no difference in resistance levels (p = 0.44 and 0.25). The ‘rabbit ears’ in **B** indicate that the medians are also the highest values. Data and details of the analysis are included in the code for Figure 5.

**Supplementary Table 1.**
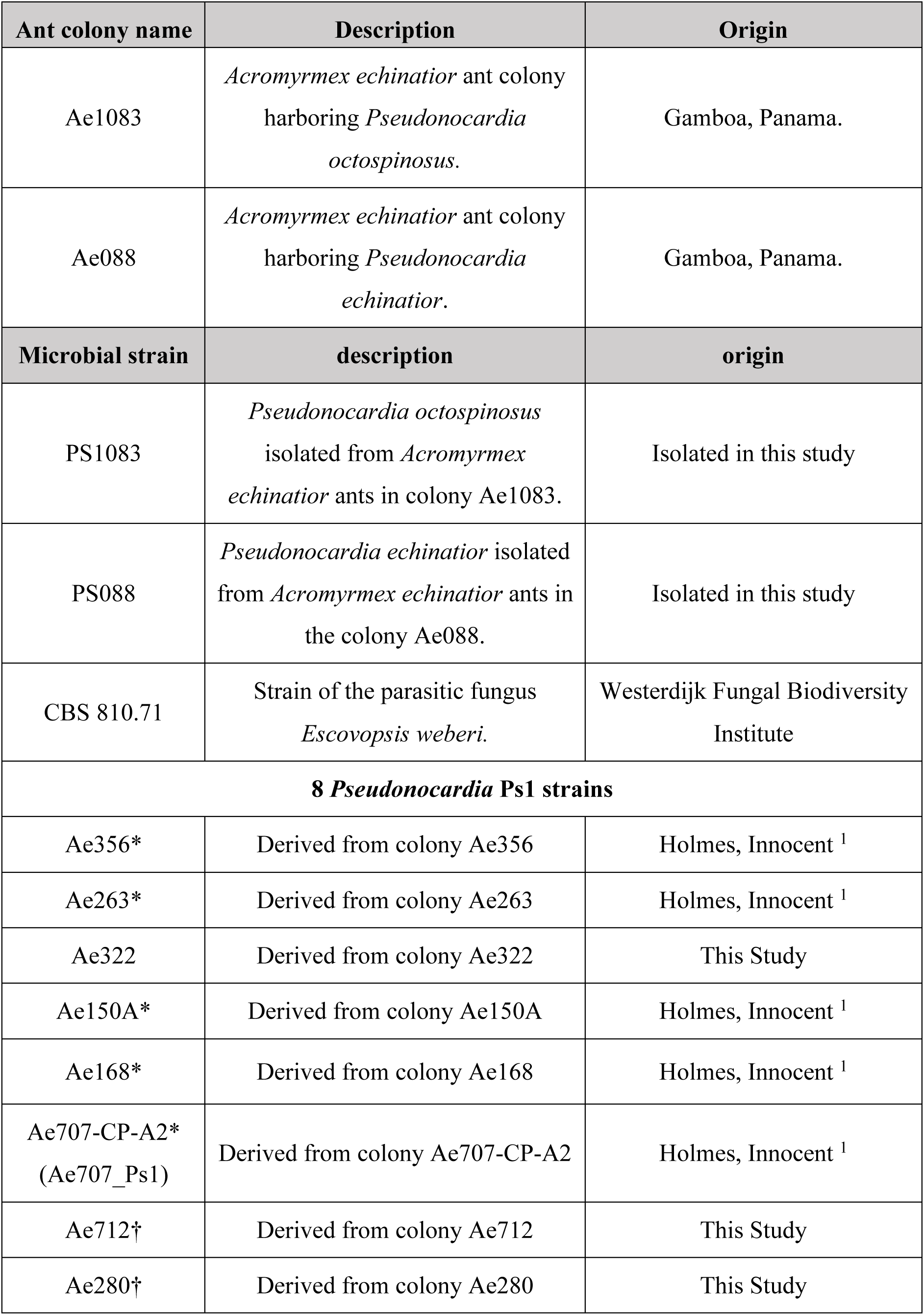
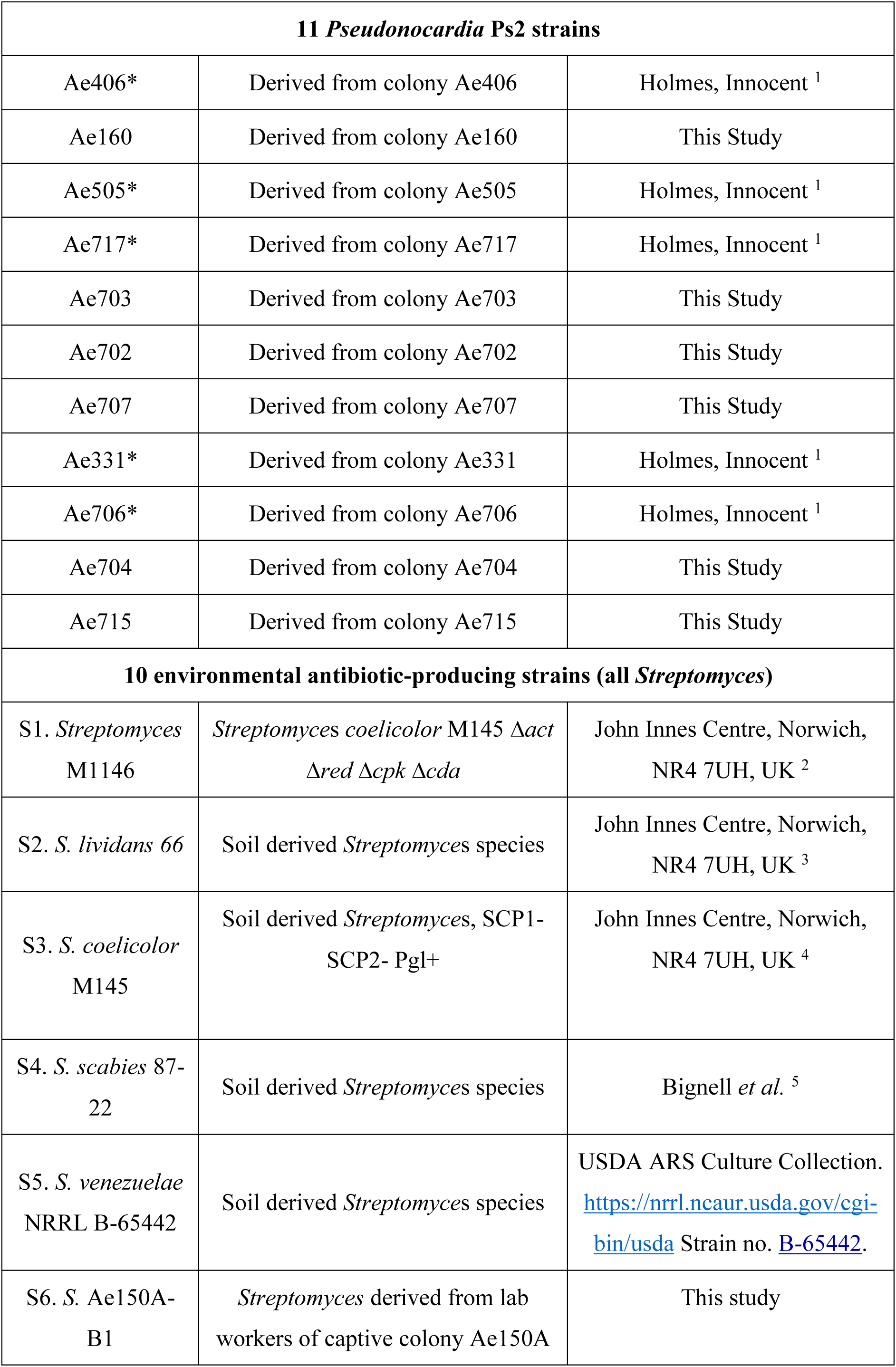
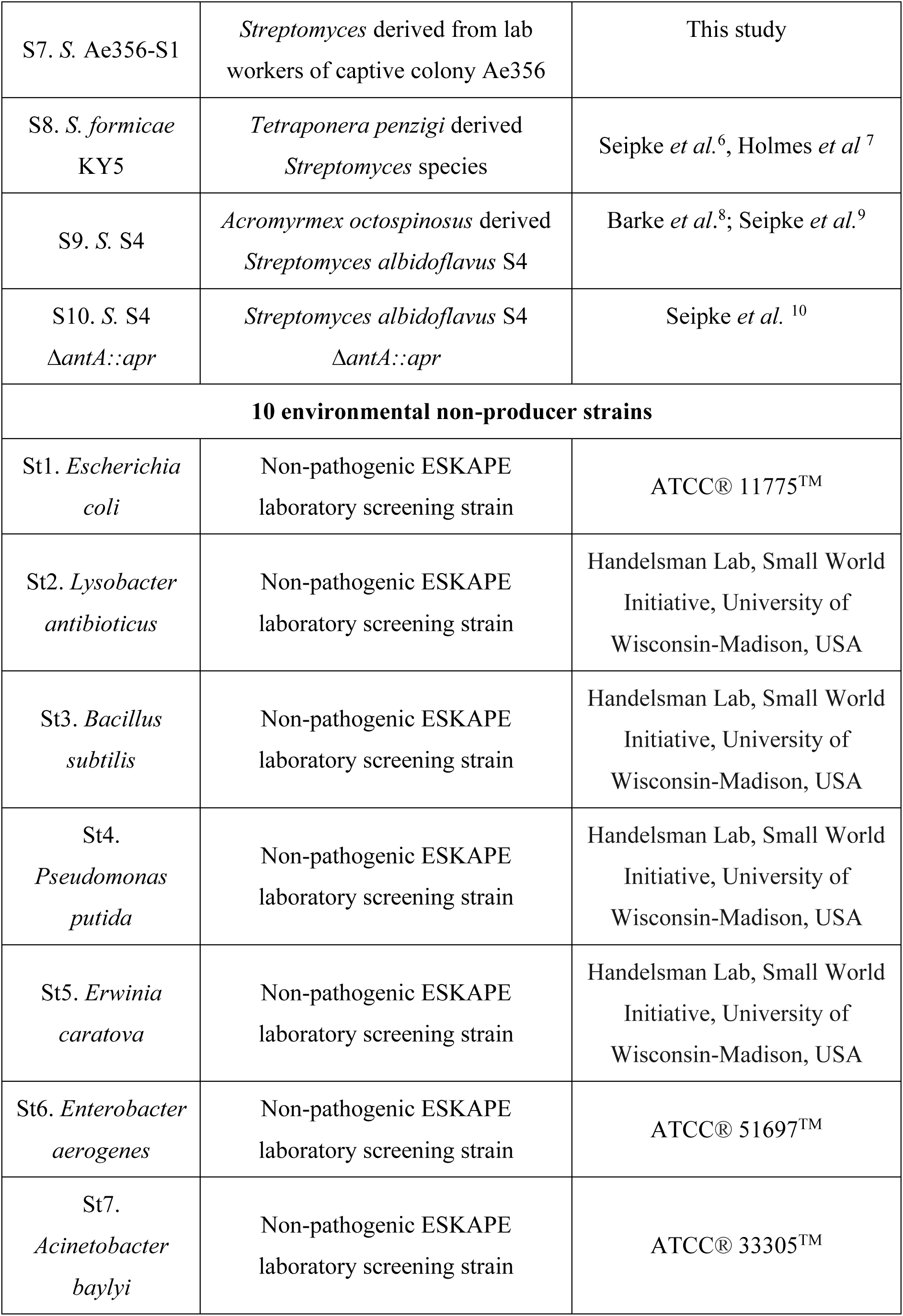
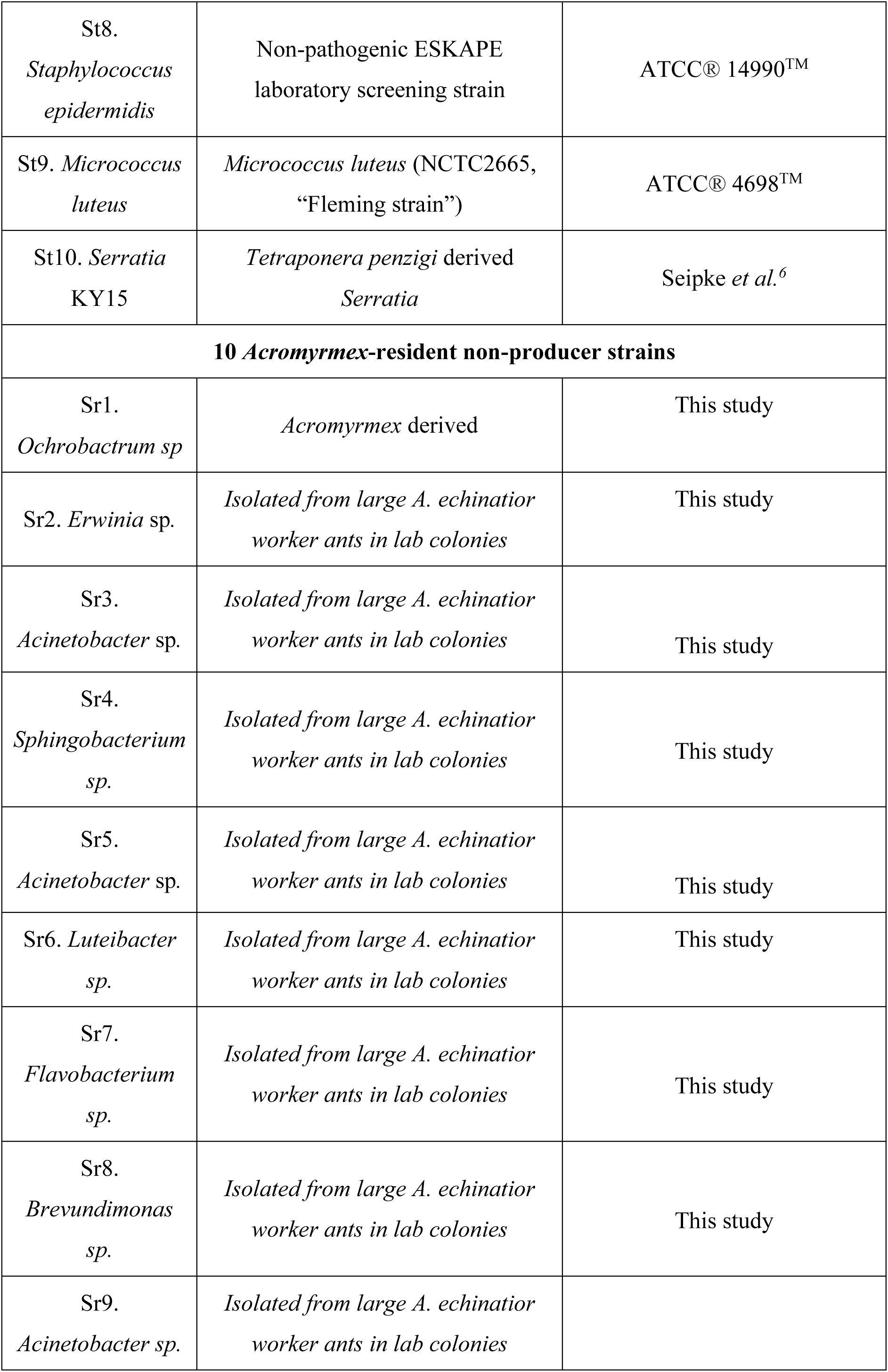
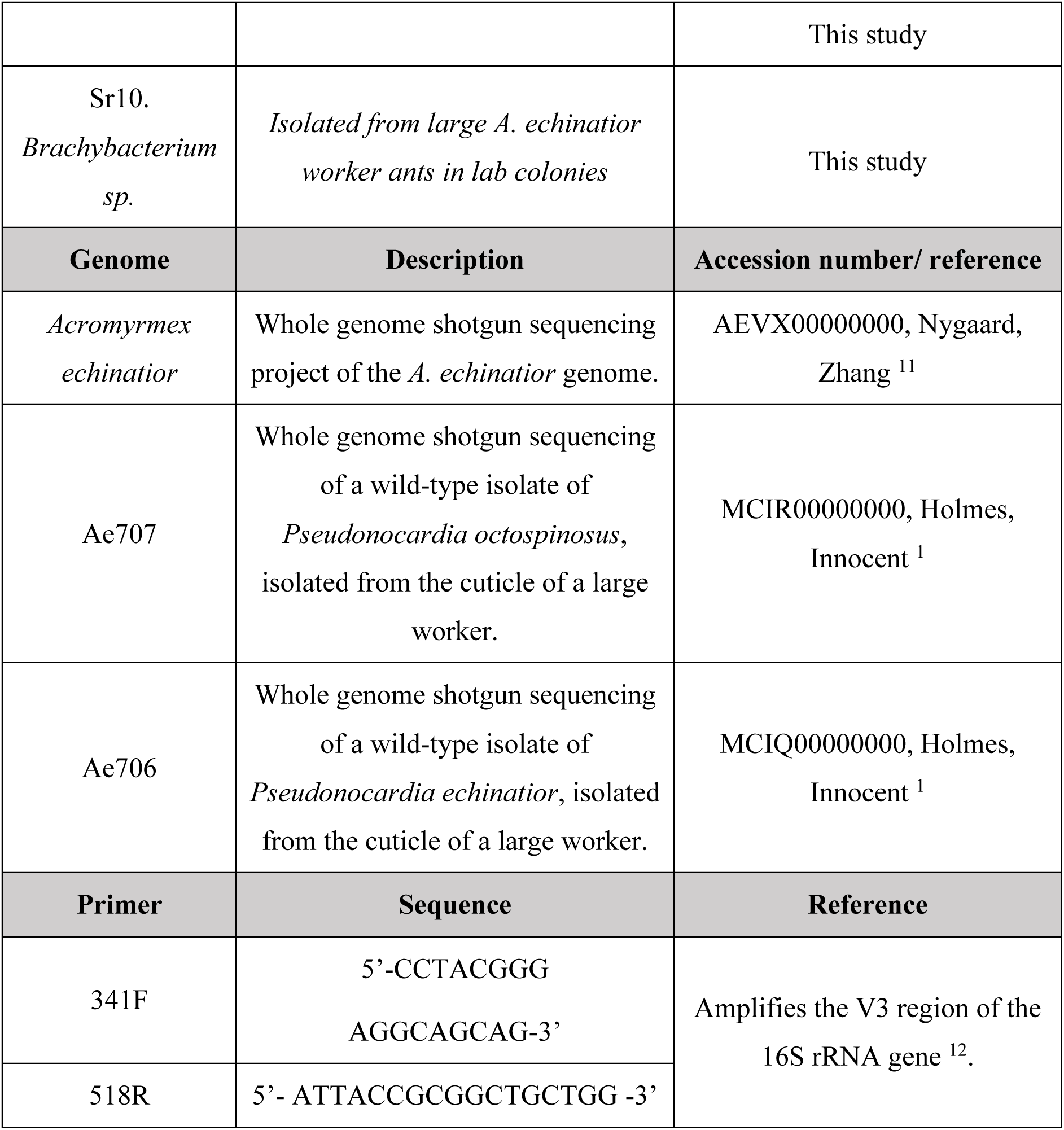
Details of ant colonies, bacterial strains and fungal strains, reference genomes and primers used in experiments. \**Pseudonocardia* strains that have been genome-sequenced. †*Pseudonocardia* strains that were only used in the growth-rate experiment with non-producers.

**Supplementary Table 2.**
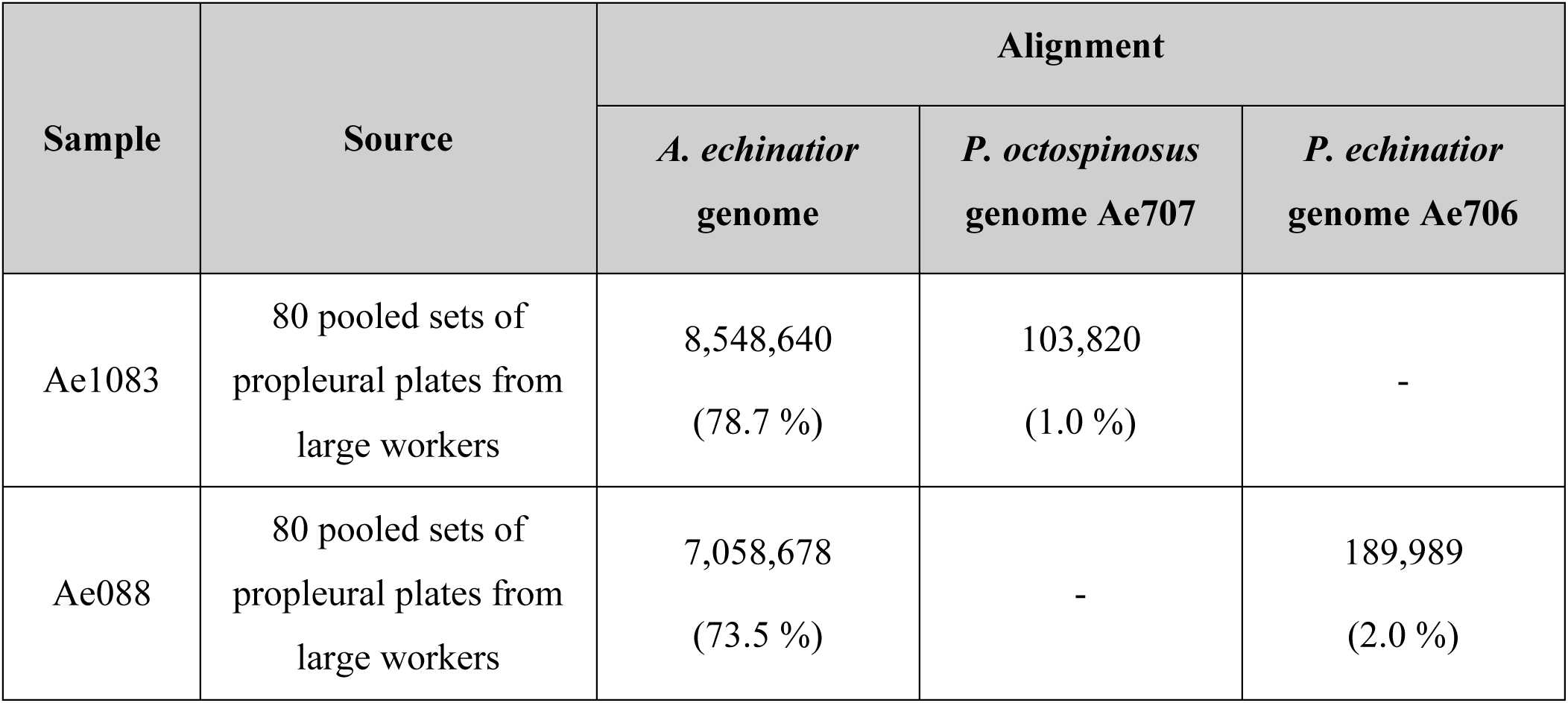
The total number of RNA-sequencing reads (and percentage of total reads in brackets) originating from propleural plate samples taken from the ant colonies Ae1083 or Ae088, respectively, that successfully aligned to the *A. echinatior* genome (Supplementary Table 1) and to the genomes of the *Pseudonocardia* species associated with the ant colony of origin (*P. octospinosus* or *P. echinatior,* respectively).

**Supplementary Table 3.**
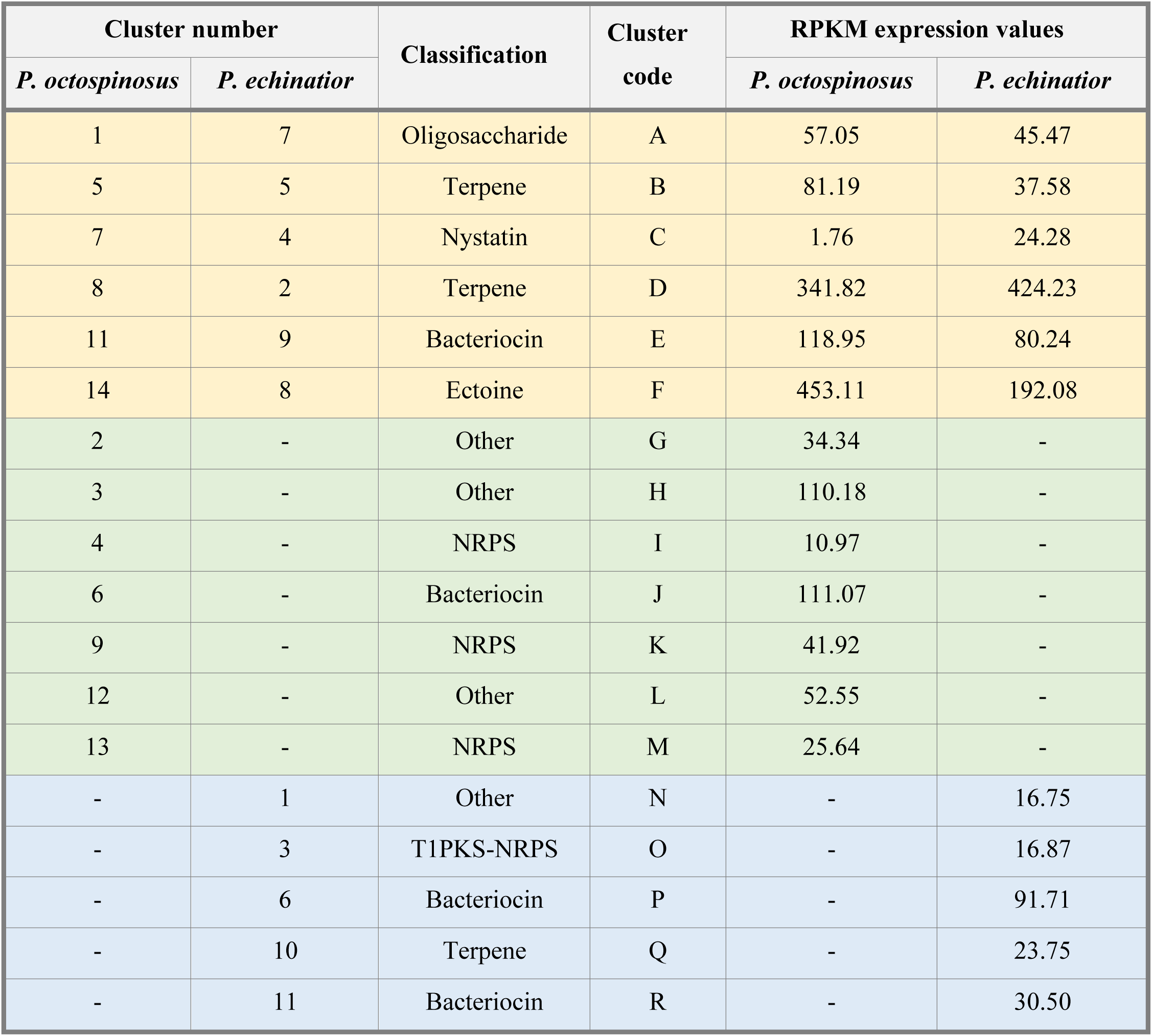
Secondary metabolite BGCs in the *Pseudonocardia* mutualist genomes (table adapted from Holmes et al.^1^) and their associated expression values (in reads per kilobase of transcript per million mapped reads, RPKM) in RNA-sequencing experiments. Yellow rows are BGCs shared between strains. Green and blue represent BGCs that are unique to *P. octospinosus* and *P. echinatior*, respectively.

**Supplementary Table 4.** Media recipes and antibiotics used in this study.

| Medium Name | Component | g L<sup>-1</sup> dH<sub>2</sub>O |  |
| --- | --- | --- | --- |
| Soya Flour Mannitol<br>(SFM) Agar | Soy flour | 20 |  |
|  | Mannitol | 20 |  |
|  | Agar | 20 |  |
| Potato Glucose Agar<br>(PGA) | PGA (Sigma Aldrich) | 39 |  |
| Glucose, Yeast, Malt<br>(GYM) Agar | Glucose | 4 |  |
|  | Yeast extract | 4 |  |
|  | Malt extract | 10 |  |
|  | CaCO<sub>3</sub> | 2 |  |
|  | Agar | 15 |  |
| Lennox Broth (LB) | Tryptone | 10 |  |
|  | NaCl | 10 |  |
|  | Yeast extract | 5 |  |
| Antibiotic | Concentrations Used | Target | Compound Type |
| Chloramphenicol | 0, 2.5, 5, 10, 25, 50<br>µg/ml | Translation<br>inhibitor | Synthetic |
| Rifampicin | 0, 0.5, 1, 2, 5, 10 µg/ml | Translation<br>inhibitor | Polyketide |
| Streptomycin | 0, 5, 10, 20, 50, 100<br>µg/ml | Translation<br>inhibitor | Aminoglycoside |
| Vancomycin | 0, 1, 2, 4, 10, 20 µg/ml | Cell wall<br>synthesis<br>inhibitor | Glycopeptide |
| Phosphomycin | 0, 5, 10, 20, 50, 100<br>µg/ml | Cell wall<br>synthesis<br>inhibitor | Small molecule |
| Nalidixic Acid | 0, 2.5, 5, 10, 25, 50<br>µg/ml | DNA gyrase<br>inhibitor | Synthetic<br>quinolone |
| Apramycin | 0, 5, 10, 20, 50, 100<br>µg/ml | Translation<br>inhibitor | Aminoglycoside |
| Ampicillin | 0, 10, 20, 40, 100, 200<br>µg/ml | Cell wall<br>synthesis<br>inhibitor | β lactam |

## Supplementary Note 1: Supplementary methods

### Isotope ratio mass spectrometry analysis

The ^13^C composition of ants fed on a ^13^C-labelled diet was determined by using a coupled Delta plus XP Isotope Ratio Mass Spectrometer/Flash HT Plus Elemental Analyser (Thermo Finnigan) in the University of East Anglia Analytical Facility. Ants were fed on a 5% glucose solution (w/v) for 10 days; five ants were fed a ^13^C glucose solution and five were fed on a ^12^C glucose solution. After 10 days, ants were washed once in 70% ETOH, then sequentially in sterile dH_2_O before drying on filter paper. The ants were then flash frozen and stored at -80°C until being placed in a ScanVac Coolsafe freeze dryer for 5 days. Each ant was then put into an individual 75 μl tin capsule (Elemental Microanalysis); capsules were loaded into an automatic sampler and completely converted to CO_2_, N_2_ and H_2_O through combustion in an excess of oxygen (oxidation was carried out at 1020°C, followed by reduction at 650°C). Nitrous oxides formed during combustion where reduced using Cu. Helium was used as a carrier gas. After passing through a water trap (MgClO_4_), the gases were separated chromatographically on an isothermal GC column (Thermo PTFE, 0.8m, 50°C); the resulting peaks sequentially entered the ion source of the Isotope Ratio Mass Spectrometer. Gas species were then measured using a Faraday cup universal collector array, with masses of 44, 45 and 46 being monitored for the analysis of CO_2_. Casein and collagen were used to calibrate the system and normalize the data post run; these standards have been calibrated against international certified standards and have an assigned δ^13^C value. Empty tin capsules were used as blanks. Each sample was analysed in triplicate. The ^13^C content of samples was reported as the ^13^C atom percent, which was calculated using the following formula:

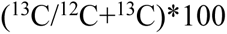

### Fluorescent microscopy of ant feeding habits

To confirm that the glucose water diet did not spread over the ant cuticle, a 20% glucose solution labelled with non-toxic fluorescent green drain tracing dye (Hydra) was fed to ants. Five mg ml^-1^ of dye was added to a 20% glucose solution, of which 300 µl was supplied to ants in the cap of an Eppendorf. Ants were sampled just after taking a feed, and following 6 and 24 hours of being exposed to the dye, to trace the spread of the solution over time. After sampling, ants were carefully fixed on their backs and brightfield and fluorescent images were acquired using a Zeiss M2 Bio Quad SV11 stereomicroscope. The samples were illuminated either with a halogen lamp (brightfield) or a 100W Hg arc lamp (fluorescence) and reflected-light images were captured with an AxioCam HRc CCD camera and AxioVision software (Carl Zeiss, Cambridge, UK). Green fluorescence was excited with light passed through a 470 nm filter (40 nm bandpass) and the emission was collected through a 525 nm filter (50 nm bandpass).

### RNA extraction protocol

A modified version of the Qiagen RNeasy Micro Kit protocol was used for all RNA extractions. Briefly, 700 μl of RLT buffer (with 1% beta mercaptoethanol) was added to each lysis matrix E tube before the samples could thaw. Tubes were then placed in a FastPrep-24™ 5G benchtop homogenizer (MP Biomedicals) and disrupted for 40 seconds at 6 m/s. Samples were then centrifuged for 2 minutes at 13’000 rpm and the supernatant was collected into a QIAshredder tube. This was centrifuged for 2 minutes at 13’000 rpm to homogenize the lysate. The resulting flow-through was mixed vigorously with 700 μl acidic phenol chloroform, then allowed to rest for 3 minutes at room temperature before centrifugation for 20 minutes at 13’000 rpm. The upper phase was then collected and a 50% volume of 96% ethanol was added. The mixture was then placed into a minelute column supplied with the Qiagen RNeasy Micro Kit. The kit protocol (including the on-column DNase I treatment) was then followed through to elution of the RNA, at which point 50 μl of RNase free water (heated to 37°C) was added to the column membrane and incubated at 37°C for 5 minutes, before centrifuging for one minute at 13’000 rpm to elute the RNA. To remove any remaining DNA, RNA was treated with the turbo DNase kit (Invitrogen): 5 μl of 10x buffer and 2 μl of Turbo DNase was added to 50 μl of RNA and incubated at 37°C for 25 minutes. RNA was then purified using the Qiagen Micro RNA Kit clean-up protocol.

### Density gradient ultracentrifugation and fractionation

Density gradient ultracentrifugation was carried out to separate ^13^C labelled (“heavy”) from un-labelled (“light”) RNA within the same RNA sample. To make one complete gradient solution for the ultracentrifugation of one RNA sample, 4.5 ml of caesium trifluoroacetate (CsTFA, ∼2 g ml-1, GE Healthcare, Munich, Germany) was added to 850 μl of gradient buffer and 197.5 μl formamide. Gradient buffer was made using an established protocol ^13^, after which 270 ng of RNA from one replicate sample was added to the gradient solution and the refractive index (R.I) of a 60 μl aliquot measured using a refractometer (Reichert Analytical Instruments, NY, USA). R.I was normalized to 1.3725 (approximately 1.79 g ml-1 CsTFA). Samples underwent ultracentrifugation in a Beckman Optima XL-100K ultracentrifuge for 50 hours at 20°C, 38’000 rpm with a vacuum applied, using a Vti 65.2 rotar (Beckman Coulter, CA, USA). Deceleration occurred without brakes. Following centrifugation, samples were divided into 12 fractions using a peristaltic pump to gradually displace the gradient, according to an established protocol ^14^. The R.I of fractions was measured to confirm the formation of a linear density gradient. RNA was precipitated from fractions by adding 1 volume of DEPC-treated sodium acetate (1M, pH 5.2), 1 μl (20 μg) glycogen (from mussels, Sigma Aldrich) and 2 volumes of ice cold 96% ETOH. Fractions were incubated over night at -20°C then centrifuged for 30 minutes at 4°C at 13’000 x g, before washing with 150 μl of ice cold 70% ETOH and centrifuging for a further 15 minutes. Pellets were then air-dried for 5 minutes and re-suspended in 15 μl of nuclease free water.

### QPCR experiments

QPCR experiments were carried out to enable the quantification of 16S rRNA gene copy number across the cDNA fractions resulting from density gradient ultracentrifugation of RNA-SIP samples. 1 μl of either template cDNA, standard DNA, or dH_2_O as a control, was added to 24 μl of reaction mix containing 12.5 μl of 2x Sybr Green Jumpstart Taq Ready-mix (Sigma Aldrich), 0.125 μl each of the primers PRM341F and 518R (Supplementary Table 1), 4 μl of 25 mM MgCl2, 0.25 μl of 20 μg μl^-1^ Bovine Serum Albumin (Sigma Aldrich), and 7 μl dH_2_O. Sample cDNA, standards (a dilution series of the target 16S rRNA gene at known quantities), and negative controls were quantified in duplicate. Reactions were run under the following conditions: 95°C for 10 mins; 40 cycles of 95°C for 15 sec, 55°C for 30 sec, and 72°C for 30 sec; plate read step at 83.5°C for 10 seconds (to avoid primer dimers); 96°C for 15 sec; 100 cycles at 55°C-95°C for 10 secs, ramping 0.5°C per cycle, followed by a plate read. Reactions were performed in 96-well plates (Bio-Rad). The threshold cycle (CT) for each sample was then converted to target molecule number by comparing to CT values of a dilution series of target DNA standards.

### Processing of reads generated from RNA sequencing experiments

The quality of Illumina sequences (returned from Vertis Biotechnologie AG) was assessed using the program FastQC (Babraham Institute, Cambridge, UK), before using TrimGalore version 0.4.5 (Babraham Institute, Cambridge, UK) to trim Illumina adaptors and low quality base calls from the 3’ end of reads (an average quality phred score of 20 was used as cut-off). After trimming, sequences shorter than 20 base pairs were discarded. Trimmed files were then aligned to the reference genome for *Acromyrmex echinatior*^11^, Supplementary Table 1, and the appropriate *Pseudonocardia* genome (either Ae707 for the sample from colony Ae1083, or Ae706 for the sample from colony Ae088^1^, see Supplementary Table 1 for genome information). All alignments were done using the splice-aware alignment program HiSat2 ^15^ with the default settings. For each cuticular sample, reads that had mapped successfully to their respective *Pseudonocardia* genomes (Supplementary Table 3) were then mapped back to the ant genome (and vice versa) to check that reads did not cross-map between the two genomes (i.e that they were either uniquely ant or bacterial reads) - reads that did not cross-map were retained for downstream analysis. Following alignment, the program HTSeq^16^ was used to count mapped reads per annotated coding sequence (CDS) using the General Feature Format (GFF) file containing the annotated gene coordinates for each reference genome. Reads that mapped to multiple locations within a genome were discarded at this point and only uniquely mapped reads were used in the counting process. Read counts per CDS were then converted to reads per kilobase of exon model per million reads (RPKM) by extracting gene lengths from the GFF file. Converting reads to RPKM values normalizes counts for RNA length and for differences in sequencing depth, which enables more accurate comparisons both within and between samples ^17^.

### Isolation of Pseudonocardia and Escovopsis bioassays

The *Pseudonocardia* strains PS1083 and PS088 (Supplementary Table 1) were isolated from the propleural plates of individual *Acromyrmex echinatior* leafcutter ants taken from colonies Ae1083 and Ae088 (Supplementary Table 1), respectively. Similarly, *Pseudonocardia* strains Ae322, Ae712, Ae280, Ae160, Ae703, Ae702, Ae707, Ae704 and Ae715 were isolated from large worker ants from colonies with the same labels maintained at the University of Copenhagen (these strains were only used for the growth-rate experiments). A sterile needle was used to scrape bacterial material off the propleural plates on the ventral part of the thorax; this was then streaked over Soya Flour Mannitol (SFM, Supplementary Table 4) agar plates and incubated at 30°C. Resulting colonies resembling *Pseudonocardia* were purified by repeatedly streaking single colonies onto SFM agar plates. Spore stocks were created using an established protocol^18^. The taxonomic identity of each *Pseudonocardia* isolate was confirmed via colony PCR and 16S rRNA sequencing, as described in Holmes *et al.*^1^. Each resulting sequence was also aligned to both the *Pseudonocardia octospinosus* and *Pseudonocardia echinatior* 16S rRNA gene sequences ^1^ to reveal their percentage identities to each of the two species. Antifungal bioassays were carried out using the *Escovopsis weberi* strain CBS 810.71, acquired from the Westerdijk Fungal Biodiversity Institute (Supplementary Table 1). *E. weberi* was actively maintained on potato glucose agar (PGA, Supplementary Table 4) at room temperature. Fungal mycelia were transferred to a fresh plate every month. For bioassays, a plug of actively growing mycelium was transferred from PGA plates to the edge of a Glucose Yeast Malt (GYM, Supplementary Table 4) agar plate with a growing *Pseudonocardia* colony, using the end of a sterile glass Pasteur pipette. Plates were then left at room temperature for 2 weeks. A zone of clearing around the *Pseudonocardia* colony indicated the presence of antifungal activity. Three replicate experiments (with three replicate bioassay plates per *Pseudonocardia* strain) were carried out, whereby different *E. weberi* starter plates were used as an inoculum.

### Individual growth-rate experiments

To create the *Pseudonocardia*-infused media, lawns of each of the 19 isolates were grown (Supplementary Table 1), plating 30 μl of spores (in 20% glycerol) onto 90 mm SFM agar plates (Supplementary Table 4). The control plates were inoculated with 20% glycerol only. We incubated these plates at 30 °C for 6 weeks, which ultimately produced confluent lawns from 17 strains that could be included in the experiments (6 Ps1, 11 Ps2). Once each plate was fully covered, we flipped the agar to reveal a surface open for colonisation. The 10 environmental producer strains and the 10 environmental non-producer strains (Supplementary Table 1) were inoculated onto the plates. Each of the plates received 10 evenly spaced colonies, with 3 replicates, generating 2 invader types x 17 Ps-media-types x 3 replicates = 102 Ps-infused plates and 2 invader types x 3 replicates = 6 control plates, for 1020 treatment and 60 control inoculations. Each strain inoculation used 5 μl of solution (approx. 1 x 106 cells per ml in 20% glycerol), spotted at evenly spaced positions and without coming into direct contact. All plates were incubated at 30 °C for five days, after which photographs were taken.

Images were processed in Fiji software^19, 20^, creating binary negatives (black & white) so automated tools could identify discrete areas of growth (black) and measure growth areas for each invading strain; in the few cases where binary image resolution was insufficient, outlines were added manually before area calculation. 48 producer-inoculated and 57 non-producer-inoculated treatment measurements were excluded because plate condition had deteriorated to become unscorable or they were contaminated, leaving final sample sizes of 1020-48-57= 915 treatment inoculations and 60 controls.

The second growth-rate experiment compared the 10 *Acromyrmex*-resident, non-producer strains with 9 of the environmental producer strains (1 of the 10 inoculations failed to grow). All 19 *Pseudonocardia* strains grew sufficiently to be included in this experiment, and each plate was again inoculated with 10 or 9 evenly spaced colonies. Starting sample sizes were therefore 2 invader types x 19 Ps-media-types x 3 replicates = 114 Ps-infused plates and 2 x 3 = 6 control plates, for 1083 treatment and 57 control inoculations. Fifty producer and 20 non-producer treatment measurements were excluded for the same reasons as above, leaving final sample sizes of 1083-50-20=1013 treatment and 57 control inoculations, scored as above.

### Pairwise competition experiment

Experiments were set up to test whether *antibiotic-producer strains* could win in direct competition against *non-producing strains,* both on normal media and on media infused with *Pseudonocardia* secondary metabolites. To create the *Pseudonocardia*-infused media, we plated 30 μl of spores (in 20% glycerol) onto 50 mm SFM agar plates (Supplementary Table 4). The control plates were inoculated with 20% (v/v) glycerol only. We incubated these plates at 30 °C for 6 weeks, which ultimately produced confluent lawns. As above, the agar was flipped to reveal a surface open for colonisation. Environmental *producers* and *non-producers* were then coinoculated onto these media (as well as on control media with no *Pseudonocardia* present), and we measured the outcome of competition as a win, loss or draw. To keep the number of tests manageable, we used two combinations of *Pseudonocardia*-infused media and *Streptomyces*: *Pseudonocardia octospinosus* (strain Ae707-CP-A2) **+** *Streptomyces* S8 and *Pseudonocardia echinatior* (strain Ae717) **+** *Streptomyces* S2 (Supplementary Table S1). We competed the two *Streptomyces* strains (S2, S8) against the 10 environmental *non-producer* strains (20 pairings). Each *Strepromyces* strain was prepared as 10^6^ spores per ml in 20% glycerol. Each *non-producer* strains was grown overnight in 10 ml of Lennox broth (Supplementary Table 4), before subculturing (1:100 dilution) into 10 ml of fresh Lennox, and incubating at 37 °C for 3-4 hours. The OD_600_ was then measured, assuming that OD_600_ = 1 represented 8 × 10^9^ cells. Similar dilutions of 10^6^ cells per ml were made for each *non-producer* strain in 20% (v/v) glycerol, after which *producer* and *non-producer* preparations were mixed at a ratio of 1:1 (v/v) and co-inoculated as a mixture of 20 µl (10^4^ spore-cells of each) on the designated *Pseudonocardia*-infused media with 5 replicates per pairing. We used 150 plates for the S8 experiment (including 100 control plates; 10 replicates per pairing) and 100 plates for the S2 experiment (including 50 control plates; 5 replicates per pairing). Plates were incubated at 30 °C for 5 days before imaging, after which images were scored with respect to the *producer* as: win (dominant growth), draw (both strains growing with no clear dominant) or lose (little or no visible growth), always with reference to images of each strain grown alone on control medium to minimise observer bias. One plate’s outcome was too ambiguous to score and was discarded. All plates were independently scored by two observers, one using photos of the original images, which produced datasets giving the same statistical results. We report the direct observer’s scores (Supplementary Methods). For analysis, draw outcomes were omitted, and a general linear mixed-effects model, including non-antibiotic-producer strain as a random intercept (10 groups), was used to test for an effect of the medium term (Control vs. Ps1/2-infused) on competitive outcome (Win vs. Loss) ((*lme4::glmer*(outcome ∼ medium + (1 | non.producer.strain), family = binomial)). Significance was estimated using term deletion.

### Antibiotic resistance assays

The key assumption of screening theory is that antibiotic-*producers* are better at resisting antibiotics, as measured by growth rates in the presence of antibiotics, because this correlation is what allows *producer* strains to better endure the demanding environment produced by *Pseudonocardia*. We tested this assumption by growing the 10 environmental *producer* strains, the 10 environmental *non-producer* strains, and the 10 resident non-producers strains (Supplementary Table 1) in the presence of 8 different antibiotics (Supplementary Table 4), representing a range of chemical classes and modes of action. Antibiotics were added to 1 ml of LB-Lennox/SFM medium (Supplementary Table 4) in a 24-well microtitre plate at 6 different concentrations. The relative concentration range was the same for each antibiotic, although actual concentrations reflected activity (Supplementary Table 4). *Producers* and *non-producers* were inoculated onto plates and incubated at 30 °C for 7 days, then photographed. Lowest Effective Concentration (LEC, lowest concentration with inhibitory effect) and Minimum Inhibitory Concentration (MIC, lowest concentration with no growth) scores were assigned on a Likert scale of 1–6, where 1 was no resistance and 6 was resistance above the concentrations tested (adapted from generalized MIC methods; reviewed by Balouiri *et al.*^21^).

### Data Analysis

R markdown-format scripts, input datafiles, and html output files for the analyses in Main Text Figures 4, 5, and 6, and in Supplementary Figure 9 are provided as a single R project folder at <u>github.com/dougwyu/Worsley_et_al_screening_test_R_code</u>

## References

1. McCutcheon JP, Boyd BM, Dale C. The Life of an Insect Endosymbiont from the Cradle to the Grave. Curr Biol 29, R485–R495 (2019).

2. Brucker RM, Bordenstein SR. The hologenomic basis of speciation: gut bacteria cause hybrid lethality in the genus Nasonia. Science 341, 667–669 (2013).

3. Rosshart SP, et al. Wild Mouse Gut Microbiota Promotes Host Fitness and Improves Disease Resistance. Cell 171, 1015–1028 e1013 (2017).

4. Sommer F, Backhed F. The gut microbiota--masters of host development and physiology. Nat Rev Microbiol 11, 227–238 (2013).

5. Maynard CL, Elson CO, Hatton RD, Weaver CT. Reciprocal interactions of the intestinal microbiota and immune system. Nature 489, 231–241 (2012).

6. Granato ET, Meiller-Legrand TA, Foster KR. The Evolution and Ecology of Bacterial Warfare. Curr Biol 29, R521–R537 (2019).

7. Foster KR, Schluter J, Coyte KZ, Rakoff-Nahoum S. The evolution of the host microbiome as an ecosystem on a leash. Nature 548, 43–51 (2017).

8. Widder S, et al. Challenges in microbial ecology: building predictive understanding of community function and dynamics. ISME J 10, 2557–2568 (2016).

9. Hoye BJ, Fenton A. Animal host-microbe interactions. J Anim Ecol 87, 315–319 (2018).

10. Archetti M, Ubeda F, Fudenberg D, Green J, Pierce NE, Yu DW. Let the right one in: a microeconomic approach to partner choice in mutualisms. Am Nat 177, 75–85 (2011).

11. Scheuring I, Yu DW. How to assemble a beneficial microbiome in three easy steps. Ecol Lett 15, 1300–1307 (2012).

12. Boza G, Worsley SF, Yu DW, Scheuring I. Efficient assembly and long-term stability of defensive microbiomes via private resources and community bistability. Plos Comput Biol 15, e1007109 (2019).

13. Heil M. Let the best one stay: screening of ant defenders by Acacia host plants functions independently of partner choice or host sanctions. Journal of Ecology 101, 684–688 (2013).

14. Tragust S, et al. Formicine ants swallow their highly acidic poison for gut microbial selection and control. eLife 9, e60287 (2020).

15. Itoh H, et al. Host-symbiont specificity determined by microbe-microbe competition in an insect gut. Proc Natl Acad Sci U S A 116, 22673–22682 (2019).

16. Ranger CM, et al. Symbiont selection via alcohol benefits fungus farming by ambrosia beetles. Proc Natl Acad Sci U S A 115, 4447–4452 (2018).

17. Sen R, Ishak HD, Estrada D, Dowd SE, Hong E, Mueller UG. Generalized antifungal activity and 454-screening of *Pseudonocardia* and *Amycolatopsis* bacteria in nests of fungus-growing ants. Proc Natl Acad Sci USA 106, 17805 (2009).

18. Andersen SB, Hansen LH, Sapountzis P, Sorensen SJ, Boomsma JJ. Specificity and stability of the *Acromyrmex*-*Pseudonocardia* symbiosis. Mol Ecol 22, 4307–4321 (2013).

19. Chapela IH, Rehner SA, Schultz TR, Mueller UG. Evolutionary History of the Symbiosis Between Fungus-Growing Ants and Their Fungi. Science 266, 1691 (1994).

20. Mueller UG, Rehner SA, Schultz TR. The evolution of agriculture in ants. Science 281, 2034–2038 (1998).

21. Currie CR. A community of ants, fungi, and bacteria: a multilateral approach to studying symbiosis. Annu Rev Microbiol 55, 357–380 (2001).

22. De Fine Licht HH, Boomsma JJ, Tunlid A. Symbiotic adaptations in the fungal cultivar of leaf-cutting ants. Nat Commun 5, 5675 (2014).

23. Sapountzis P, Zhukova M, Shik JZ, Schiott M, Boomsma JJ. Reconstructing the functions of endosymbiotic Mollicutes in fungus-growing ants. Elife 7, (2018).

24. Reynolds HT, Currie CR. Pathogenicity of Escovopsis weberi: The parasite of the attine ant-microbe symbiosis directly consumes the ant-cultivated fungus. Mycologia 96, 955–959 (2004).

25. Heine D, et al. Chemical warfare between leafcutter ant symbionts and a co-evolved pathogen. Nat Commun 9, 2208 (2018).

26. Worsley SF, et al. Symbiotic partnerships and their chemical interactions in the leafcutter ants (Hymenoptera: Formicidae). Myrmecological News 27, 59–74 (2018).

27. de Man TJ, et al. Small genome of the fungus *Escovopsis weberi*, a specialized disease agent of ant agriculture. Proc Natl Acad Sci U S A 113, 3567–3572 (2016).

28. Currie CR, Stuart AE. Weeding and grooming of pathogens in agriculture by ants. Proc Biol Sci 268, 1033–1039 (2001).

29. Fernandez-Marin H, et al. Functional role of phenylacetic acid from metapleural gland secretions in controlling fungal pathogens in evolutionarily derived leaf-cutting ants. Proc Biol Sci 282, 20150212 (2015).

30. Fernandez-Marin H, Zimmerman JK, Rehner SA, Wcislo WT. Active use of the metapleural glands by ants in controlling fungal infection. Proc Biol Sci 273, 1689–1695 (2006).

31. Holmes NA, et al. Genome analysis of two *Pseudonocardia* phylotypes associated with *Acromyrmex* leafcutter ants reveals their biosynthetic potential. Front Microbiol 7, 2073 (2016).

32. Poulsen M, Cafaro M, Boomsma JJ, Currie CR. Specificity of the mutualistic association between actinomycete bacteria and two sympatric species of *Acromyrmex* leaf-cutting ants. Mol Ecol 14, 3597–3604 (2005).

33. Currie CR, Scott JA, Summerbell RC, Malloch D. Fungus-growing ants use antibiotic-producing bacteria to control garden parasites. Nature 398, 701–704 (1999).

34. Marsh SE, Poulsen M, Pinto-Tomás A, Currie CR. Interaction between workers during a short time window is required for bacterial symbiont transmission in *Acromyrmex* leaf-cutting ants. PloS one 9, e103269 (2014).

35. Poulsen M, Bot ANM, Currie CR, Nielsen MG, Boomsma JJ. Within-colony transmission and the cost of a mutualistic bacterium in the leaf-cutting ant *Acromyrmex octospinosus*. Funct Ecol 17, 260–269 (2003).

36. Kost C, Lakatos T, Böttcher I, Arendholz W-R, Redenbach M, Wirth R. Non-specific association between filamentous bacteria and fungus-growing ants. Naturwissenschaften 94, 821–828 (2007).

37. Barke J, et al. A mixed community of actinomycetes produce multiple antibiotics for the fungus farming ant *Acromyrmex octospinosus*. BMC Biol 8, (2010).

38. Haeder S, Wirth R, Herz H, Spiteller D. Candicidin-producing *Streptomyces* support leaf-cutting ants to protect their fungus garden against the pathogenic fungus *Escovopsis*. Proc Natl Acad Sci USA 106, 4742–4746 (2009).

39. Schoenian I, Spiteller M, Ghaste M, Wirth R, Herz H, Spiteller D. Chemical basis of the synergism and antagonism in microbial communities in the nests of leaf-cutting ants. Proc Natl Acad Sci U S A 108, 1955–1960 (2011).

40. Currie CR, Poulsen M, Mendenhall J, Boomsma JJ, Billen J. Coevolved crypts and exocrine glands support mutualistic bacteria in fungus-growing ants. Science 311, 81–83 (2006).

41. Dumont MG, Murrell JC. Stable isotope probing - linking microbial identity to function. Nat Rev Microbiol 3, 499–504 (2005).

42. Neufeld JD, Wagner M, Murrell JC. Who eats what, where and when? Isotope-labelling experiments are coming of age. ISME J 1, 103–110 (2007).

43. Whiteley AS, Thomson B, Lueders T, Manefield M. RNA stable-isotope probing. Nat Protoc 2, 838–844 (2007).

44. Andersen SB, Boye M, Nash DR, Boomsma JJ. Dynamic *Wolbachia* prevalence in *Acromyrmex* leaf-cutting ants: potential for a nutritional symbiosis. J Evol Biol 25, 1340–1350 (2012).

45. Sapountzis P, Nash DR, Schiott M, Boomsma JJ. The evolution of abdominal microbiomes in fungus-growing ants. Mol Ecol 28, 879–899 (2019).

46. Seipke RF, et al. A single *Streptomyces* symbiont makes multiple antifungals to support the fungus farming ant *Acromyrmex octospinosus*. PLoS One 6, e22028 (2011).

47. Westermann AJ, et al. Dual RNA-seq unveils noncoding RNA functions in host-pathogen interactions. Nature 529, 496–501 (2016).

48. Nygaard S, et al. The genome of the leaf-cutting ant *Acromyrmex echinatior* suggests key adaptations to advanced social life and fungus farming. Genome Res 21, 1339–1348 (2011).

49. Gershenzon J, Dudareva N. The function of terpene natural products in the natural world. Nat Chem Biol 3, 408–414 (2007).

50. Richter TK, Hughes CC, Moore BS. Sioxanthin, a novel glycosylated carotenoid, reveals an unusual subclustered biosynthetic pathway. Environ Microbiol 17, 2158–2171 (2015).

51. Nupur LN, Vats A, Dhanda SK, Raghava GP, Pinnaka AK, Kumar A. ProCarDB: a database of bacterial carotenoids. BMC Microbiol 16, 96 (2016).

52. Chikindas ML, Weeks R, Drider D, Chistyakov VA, Dicks LM. Functions and emerging applications of bacteriocins. Curr Opin Biotechnol 49, 23–28 (2018).

53. Cotter PD, Ross RP, Hill C. Bacteriocins - a viable alternative to antibiotics? Nat Rev Microbiol 11, 95–105 (2013).

54. Riley MA, Gordon DM. The ecological role of bacteriocins in bacterial competition. Trends Microbiol 7, 129–133 (1999).

55. Algburi A, Zehm S, Netrebov V, Bren AB, Chistyakov V, Chikindas ML. Subtilosin prevents biofilm formation by inhibiting bacterial quorum sensing. Probiotics Antimicrob Proteins 9, 81–90 (2017).

56. Bates D, Mächler M, Bolker B, Walker S. Fitting Linear Mixed-Effects Models Using lme4. Journal of Statistical Software; Vol 1, Issue 1 (2015), (2015).

57. Balouiri M, Sadiki M, Ibnsouda SK. Methods for in vitro evaluating antimicrobial activity: A review. J Pharm Anal 6, 71–79 (2016).

58. Currie C, Bot A, Boomsma JJ. Experimental evidence of a tripartite mutualism: bacteria protect ant fungus gardens from specialized parasites. Oikos 101, 91–102 (2003).

59. Andersen SB, Yek SH, Nash DR, Boomsma JJ. Interaction specificity between leaf-cutting ants and vertically transmitted *Pseudonocardia* bacteria. BMC Evol Biol 15, 1 (2015).

60. Zivkovic AM, German JB, Lebrilla CB, Mills DA. Human milk glycobiome and its impact on the infant gastrointestinal microbiota. Proc Natl Acad Sci U S A 108 **Suppl 1**, 4653–4658 (2011).

61. Carlstrom CI, Field CM, Bortfeld-Miller M, Muller B, Sunagawa S, Vorholt JA. Synthetic microbiota reveal priority effects and keystone strains in the Arabidopsis phyllosphere. Nat Ecol Evol 3, 1445–1454 (2019).

62. Cai T, et al. Host legume-exuded antimetabolites optimize the symbiotic rhizosphere. Mol Microbiol 73, 507–517 (2009).

63. Neal AL, Ahmad S, Gordon-Weeks R, Ton J. Benzoxazinoids in root exudates of maize attract *Pseudomonas putida* to the rhizosphere. PLoS One 7, e35498 (2012).

64. Ryan PR, Dessaux Y, Thomashow LS, Weller DM. Rhizosphere engineering and management for sustainable agriculture. Plant and Soil 321, 363–383 (2009).

65. Dowd SE, et al. Evaluation of the bacterial diversity in the feces of cattle using 16S rDNA bacterial tag-encoded FLX amplicon pyrosequencing (bTEFAP). BMC Microbiol 8, 125 (2008).

66. Dowd SE, Wolcott RD, Sun Y, McKeehan T, Smith E, Rhoads D. Polymicrobial nature of chronic diabetic foot ulcer biofilm infections determined using bacterial tag encoded FLX amplicon pyrosequencing (bTEFAP). PLoS One 3, e3326 (2008).

67. DeSantis TZ, et al. Greengenes, a chimera-checked 16S rRNA gene database and workbench compatible with ARB. Appl Environ Microbiol 72, 5069–5072 (2006).

68. R Core Team. R: A language and environment for statistical computing. In: R Foundation for Statistical Computing, Vienna, Austria. https://www.R-project.org/) (2017).

69. Kanehisa M, Sato Y, Morishima K. BlastKOALA and GhostKOALA: KEGG tools for functional characterization of genome and metagenome sequences. J Mol Biol 428, 726–731 (2016).

70. Blin K, et al. AntiSMASH 4.0-improvements in chemistry prediction and gene cluster boundary identification. Nucleic Acids Res 45, W36–W41 (2017).

## References

1. Holmes NA, et al. Genome analysis of two *Pseudonocardia* phylotypes associated with *Acromyrmex* leafcutter ants reveals their biosynthetic potential. Front Microbiol 7, 2073 (2016).

2. Gomez-Escribano JP, Bibb MJ. Engineering Streptomyces coelicolor for heterologous expression of secondary metabolite gene clusters. Microb Biotechnol 4, 207–215 (2011).

3. Hopwood DA, Kieser T, Wright HM, Bibb MJ. Plasmids, Recombination and Chromosome Mapping in Streptomyces lividans 66. Microbiology 129, 2257–2269 (1983).

4. Bentley SD, et al. Complete genome sequence of the model actinomycete *Streptomyces coelicolor* A3(2). Nature 417, 141–147 (2002).

5. Bignell DR, Seipke RF, Huguet-Tapia JC, Chambers AH, Parry RJ, Loria R. *Streptomyces scabies* 87-22 contains a coronafacic acid-like biosynthetic cluster that contributes to plant-microbe interactions. Mol Plant Microbe Interact 23, 161–175 (2010).

6. Seipke RF, Barke J, Heavens D, Yu DW, Hutchings MI. Analysis of the bacterial communities associated with two ant-plant symbioses. Microbiologyopen 2, 276–283 (2013).

7. Holmes NA, Devine R, Qin Z, Seipke RF, Wilkinson B, Hutchings MI. Complete genome sequence of *Streptomyces formicae* KY5, the formicamycin producer. J Biotechnol 265, 116–118 (2018).

8. Barke J, et al. A mixed community of actinomycetes produce multiple antibiotics for the fungus farming ant *Acromyrmex octospinosus*. BMC Biol 8, (2010).

9. Seipke RF, et al. A single *Streptomyces* symbiont makes multiple antifungals to support the fungus farming ant *Acromyrmex octospinosus*. PLoS One 6, e22028 (2011).

10. Seipke RF, Patrick E, Hutchings MI. Regulation of antimycin biosynthesis by the orphan ECF RNA polymerase sigma factor sigma (AntA.). PeerJ 2, e253 (2014).

11. Nygaard S, et al. The genome of the leaf-cutting ant *Acromyrmex echinatior* suggests key adaptations to advanced social life and fungus farming. Genome Res 21, 1339–1348 (2011).

12. Muyzer G, de Waal EC, Uitterlinden AG. Profiling of complex microbial populations by denaturing gradient gel electrophoresis analysis of polymerase chain reaction-amplified genes coding for 16S rRNA. Appl Environ Microbiol 59, 695–700 (1993).

13. Lueders T, Manefield M, Friedrich MW. Enhanced sensitivity of DNA- and rRNA-based stable isotope probing by fractionation and quantitative analysis of isopycnic centrifugation gradients. Environ Microbiol 6, 73–78 (2004).

14. Whiteley AS, Thomson B, Lueders T, Manefield M. RNA stable-isotope probing. Nat Protoc 2, 838–844 (2007).

15. Kim D, Langmead B, Salzberg SL. HISAT: a fast spliced aligner with low memory requirements. Nat Methods 12, 357–360 (2015).

16. Anders S, Pyl PT, Huber W. HTSeq- a Python framework to work with high-throughput sequencing data. Bioinformatics 31, 166–169 (2015).

17. Mortazavi A, Williams BA, McCue K, Schaeffer L, Wold B. Mapping and quantifying mammalian transcriptomes by RNA-Seq. Nat Methods 5, 621–628 (2008).

18. Kieser T, Bibb MJ, Buttner MJ, Chater KF, Hopwood DA. Practical Streptomyces Genetics. John Innes Foundation (2000).

19. Rueden CT, et al. ImageJ2: ImageJ for the next generation of scientific image data. BMC Bioinformatics 18, 529 (2017).

20. Schindelin J, et al. Fiji: an open-source platform for biological-image analysis. Nat Methods 9, 676–682 (2012).

21. Balouiri M, Sadiki M, Ibnsouda SK. Methods for in vitro evaluating antimicrobial activity: A review. J Pharm Anal 6, 71–79 (2016).

